# E2 ubiquitin conjugase Bendless is essential for PINK1-Park activity to regulate Mitofusin under mitochondrial stress

**DOI:** 10.1101/2022.10.24.513457

**Authors:** Rajit Narayanan Cheramangalam, Tarana Anand, Priyanka Pandey, Deepa Balasubramanian, Reshmi Varghese, Neha Singhal, Sonal Nagarkar Jaiswal, Manish Jaiswal

**Affiliations:** Tata Institute of Fundamental Research, Hyderabad, India; CSIR–Centre For Cellular and Molecular Biology, Hyderabad, India

**Author notes:** Contributed equally to this work (Rajit Narayanan Cheramangalam, Tarana Anand).

**Keywords:** Mitochondrial quality control, Mitophagy, Parkinsons, Leigh syndrome, LRPPRC

## Abstract

Cells under mitochondrial stress often co-opt mechanisms to maintain energy homeostasis, mitochondrial quality control and cell survival. A mechanistic understanding of such responses is crucial for further insight into mitochondrial biology and diseases. Through an unbiased genetic screen in *Drosophila*, we identify that mutations in *lrpprc2*, a homolog of the human *LRPPRC* gene that is linked to the French-Canadian Leigh syndrome, results in PINK1-Park activation. While the PINK1-Park pathway is well known to induce mitophagy, we show that in the case of *lrpprc2* mutants, PINK1-Park regulates mitochondrial dynamics by inducing degradation of the mitochondrial fusion protein Mitofusin/Marf. We also discover that Bendless, a K63-linked E2 conjugase, is a regulator of Marf, as loss of *bendless* results in increased Marf levels. We show that Bendless is required for PINK1 stability, and subsequently for PINK1-Park mediated Marf degradation under physiological conditions, and in response to mitochondrial stress as seen in *lrpprc2*. Additionally, we show that loss of Bendless in *lrpprc2* mutant eye results in photoreceptor degeneration, indicating a neuroprotective role for Bendless-PINK1-Park mediated Marf degradation. Based on our observations, we propose that certain forms of mitochondrial stress activate Bendless-PINK1-Park to limit mitochondrial fusion, which is a cell-protective response.

## Introduction

Mitochondria are dynamic organelles and their size varies in response to various cellular cues such as developmental signaling (1), metabolic needs (2) or toxin induced stress (2). This change in mitochondrial size is crucial for cellular adaptation under different physiological conditions. For example, upon amino acid deprivation, mitochondria undergo fusion, which results in increased ATP production (3) and protects mitochondria from autophagy (4). Changes in mitochondrial size requires regulation of GTPases essential for mitochondrial dynamics. While the Dynamin 1-like (DNM1/Drp1) protein mediates fission, Mitofusins (Mfn1 and Mfn2 in mammals, Marf in *Drosophila*) and Optic Atrophy 1 (OPA1) mediate the fusion of mitochondrial outer and inner membranes respectively. Several post-translational modifications, such as phosphorylation, acetylation and ubiquitination are crucial for the activity of these proteins, and thereby play an important role in determining mitochondrial size (5,6). Misregulation of these proteins, and consequently, mitochondrial dynamics, is associated with metabolic and neurodegenerative diseases (7).

The E3 ubiquitin ligase Parkin (Park in *Drosophila*, PARK2 in humans) and the kinase PINK1, which are linked to autosomal recessive early-onset Parkinsonism, are known to regulate mitochondrial quality control (8). Studies in human cancer cell lines have shown that dissipation of the mitochondrial membrane potential (MMP) can stabilize PINK1 on the outer mitochondrial membrane (OMM) leading to Park recruitment, polyubiquitination of OMM proteins and mitophagy (9–12). Several *in vivo* studies have also shown a conserved role for PINK1-Parkin in mitophagy (13–20). While PINK1-Park mediated mitophagy has been extensively studied in cells, how the PINK1-Park pathway is activated under physiological conditions *in vivo* remains elusive (21). Additionally, *in vivo* studies suggest a pro-fission role of PINK1-Park (22–26), perhaps through the regulation of mitofusin levels (27). As most of these studies utilize *PINK1* and *PARK2* mutants to study defects in mitochondrial dynamics, the mechanism by which they are regulated in vivo under various physiological conditions remains unresolved. Additionally, it is unclear as to how the PINK1-Park pathway may activate mitophagy, alter mitochondrial dynamics or selectively target certain OMM proteins in response to mitochondrial stress.

To study the regulation of Mitofusin/Marf *in vivo*, we undertook an unbiased genetic screen in *Drosophila*. From this genetic screen, we discovered that mutations in *lrpprc2 (ppr)*, a homolog of human *LRPPRC* that is required for mitochondrial mRNA stability and translation (28,29), results in activation of the PINK1-Park pathway. This activation then leads to proteasome-mediated Marf degradation, but not mitophagy. We also discovered that mutations in *bendless* (*ben*), which encodes a K63-linked E2 ubiquitin conjugase, is essential for Marf degradation in *ppr* mutants. Further, we demonstrate an essential role for Ben in regulating the stability of PINK1, which in turn is required for maintaining steady state Marf levels in healthy cells. Finally, we show that in *ppr* mutants, Ben suppresses excessive mitochondrial fusion and prevents neuronal death under mitochondrial stress.

## Results

### Loss of *ppr* results in reduced Marf levels

To identify novel regulators of mitochondrial dynamics, we screened a collection of *Drosophila* X-chromosome lethal mutations (30,31). This collection was generated to identify mutants with neurodegenerative phenotypes and has previously uncovered mutations in *Marf (32)* and several other genes required for mitochondrial function (29,33). We tested these mutants for misregulation of Marf protein using an HA-tagged Marf genomic construct (*Marf::HA*). We used the *FLP*-FRT mediated mitotic recombination strategy to create mutant clones (non-GFP cells) in a heterozygous background (GFP expressing cells) in the developing wing disc epithelium (34). This allowed us to compare Marf levels in mutant and wild type cells within the same tissue (Figure S1A-A’).

From this screen, we found that mutant clones of two independent *ppr alleles* (*ppr*^*A*^ and *ppr*^*E*^) show reduced Marf:HA levels compared to the surrounding wild type cells (Figure 1A-A’’, 1E, Figure 1B-B’). To further confirm these results, we used an independent Marf genomic rescue line, Marf::mCherry and found reduced Marf::mCherry staining in *ppr*^*A*^ mutant clones (Figure S1C-C’). To test for the possibility that the reduction in Marf::HA or Marf::mCherry is caused by reduced mitochondrial content, we checked the levels of an OMM protein Tom20 using an endogenous tagged line (Tom20::mCherry). We did not observe any change in Tom20::mCherry staining in *ppr*^*A*^ mutant clones (Figure 1B-B’’, 1E). Taken together, these data suggest that downregulation of Marf in *ppr* mutants is not due to reduced mitochondrial content. Additionally, we also looked at other proteins involved in mitochondrial dynamics — Opa1 and Drp1— using genomic tags. While we found the levels of Opa1::HA to be slightly increased in *ppr*^*A*^ mutant clones (Figure 1C-C”, 1E), Drp1::HA levels remained unaltered (Figure 1D-D”, 1E). As mutations in *ppr/LRPPRC* result in mitochondrial defects due to reduced stability of mtRNA (28,29,37), reduced Marf levels in *ppr* mutants appears to be an adaptation to segregate defective mitochondria by suppressing their fusion.

**Figure 1.**
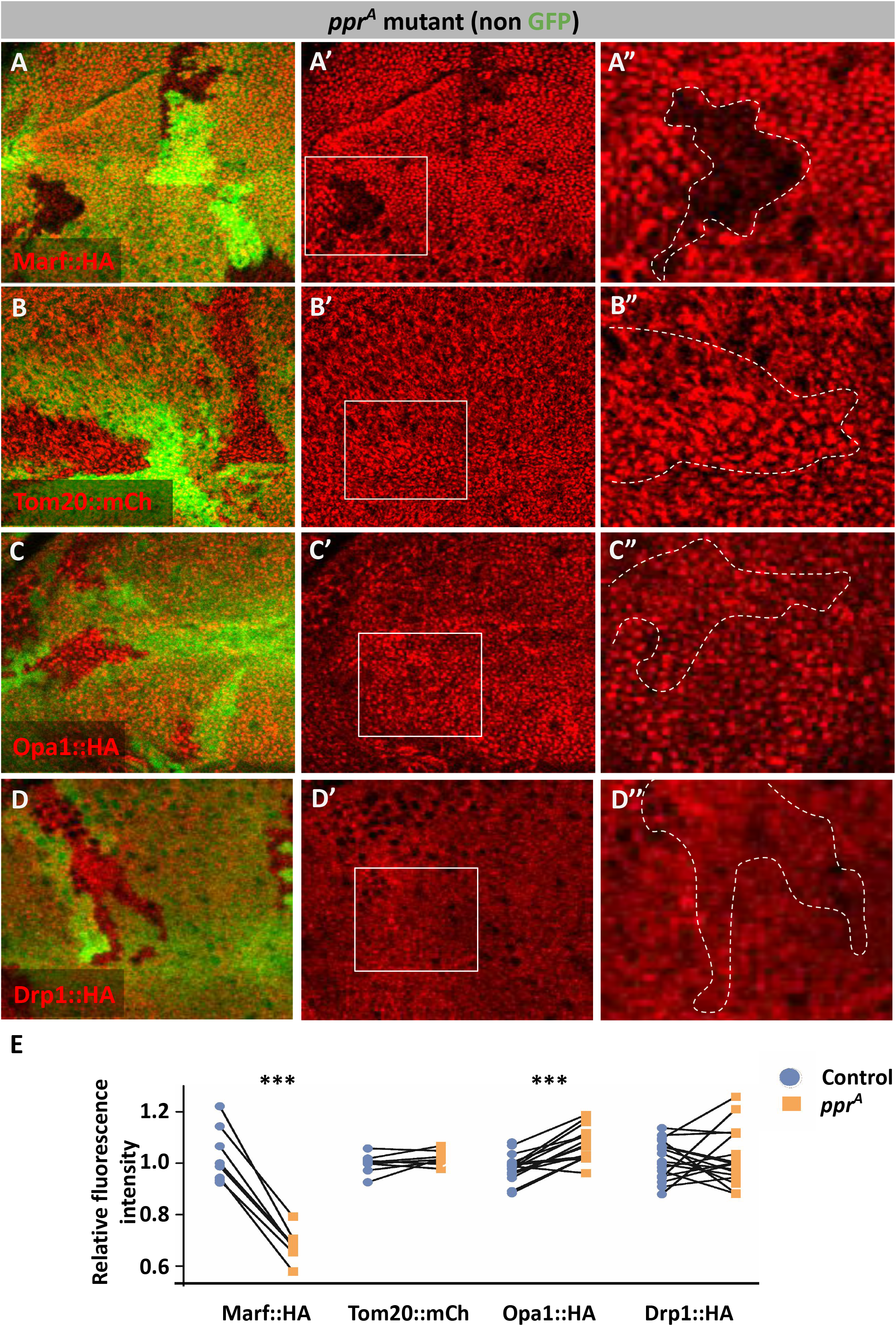
*ppr* mutants show Marf down regulation: *(A-D*”*)ppr*^*A*^ mutant clones (non green cells, *A-D* and dashed white line, *A*”*-D*”), wing discs immunostained for Marf::HA (red, *A-A*”), Tom20::mCh (red, *B-B*”), Opa1::HA (red, *C-C*”) and Drp1::HA (red, *D-D*”) (genomic rescue tags). *A*”*-D*”are magnified images of insets shown in *A*’*-D*’. *(E)*Quantification for relative fluorescence intensities of Marf::HA (n=9), Tom20::mCh (n=15), Opa1::HA (n=16) and Drp1::HA (n=19) in *ppr*^*A*^ mutant clones. Graphs represent average intensity values normalized to that of control cells. Two tailed paired t-test between control and *ppr*^*A*^ mutant cells. Significance represented by p<0.001***.

Since reduced Marf is expected to suppress mitochondrial fusion, we sought to test mitochondrial morphology in *ppr*^*A*^ mutant clones. The cells in wing discs are very compact, and hence it is difficult to study mitochondrial morphology. Hence, we created mutant clones in the peripodial membrane, which is a squamous epithelium overlying wing discs. We used anti-Complex-V staining to mark mitochondria. Interestingly, we found that mitochondrial size is increased in these *ppr*^*A*^ mutant clones (Figure S1E-E”, 1F). Similar increase in mitochondrial size has been observed in *LRPPRC* knockdown in mouse liver (35) and in C.elegans (36). As many studies have shown that mitochondrial stress can result in increased mitochondrial size (3,4,36,38), we suspect a similar mechanism results in increased mitochondrial size in *ppr* mutant cells, while an independent mitochondrial quality control mechanism may suppress their fusion by inducing Marf turnover.

### UPS dependent Marf degradation in *ppr* mutants

Reduced Marf levels in *ppr* mutant clones could be increased protein turnover via selective autophagy or ubiquitin-proteasomal system (UPS). We tested the possibility of autophagic degradation of Marf. We fed chloroquine, an inhibitor of autophagosome-lysosome fusion (39), to larvae and found that Marf::HA levels remain reduced in *ppr*^*A*^ clones (Figure 2B-B’, 2E). Moreover, we found that the levels of p62, a protein degraded primarily via autophagy, was not altered in *ppr*^*A*^ clones (Figure S1D-D’). Thus we conclude that autophagy is neither enhanced nor likely the cause of Marf reduction in *ppr* mutant clones.

**Figure 2.**
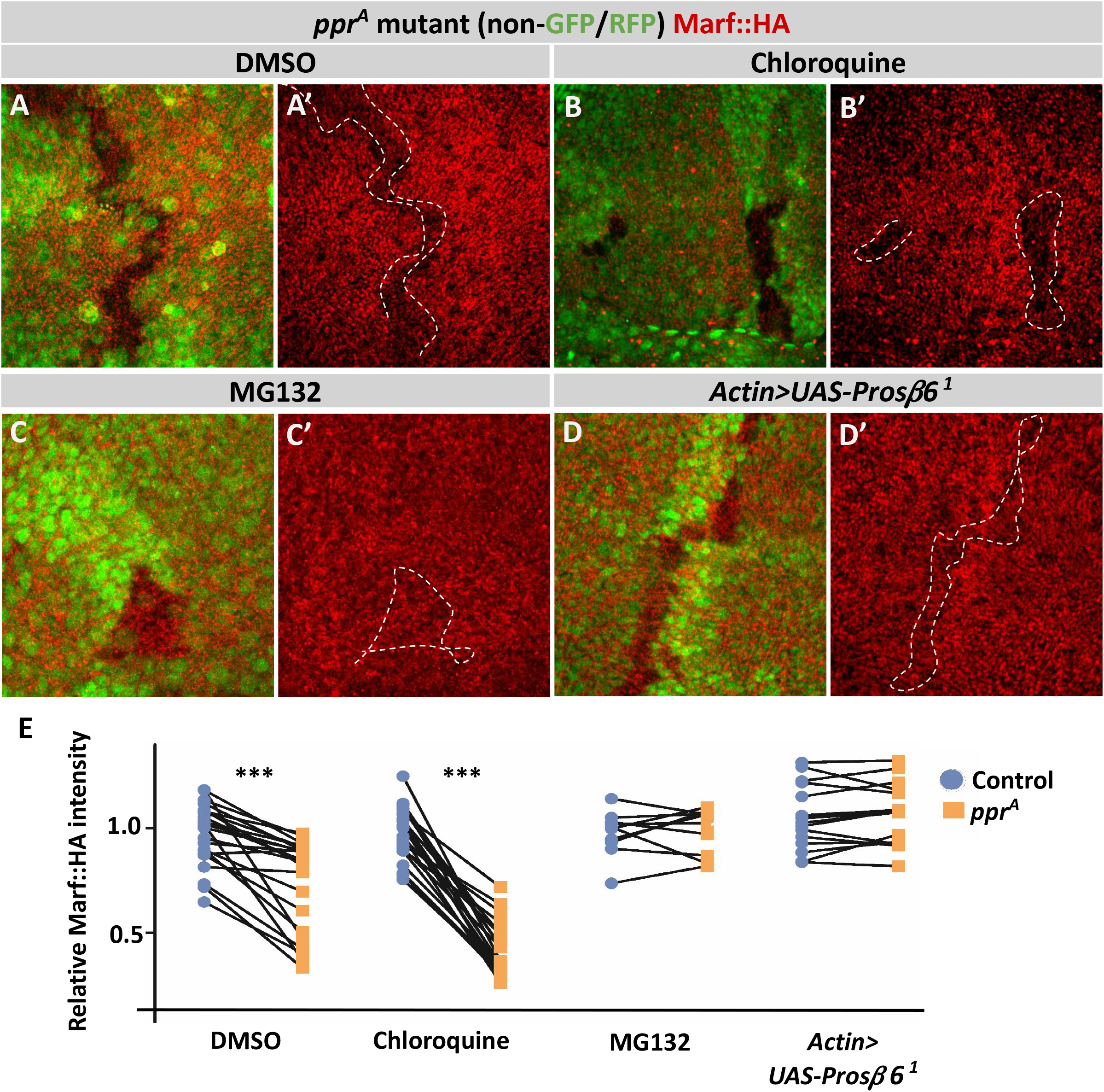
*ppr* mutants show UPS mediated Marf degradation: *(A-C*’*)ppr*^*A*^ mutant clones (non green cells, *A-C* and dashed white line, *A*’*-C*’), wing discs immunostained for Marf::HA (red) after feeding larvae with DMSO *(A-A*’), chloroquine *(B-B*’) or MG132 *(C-C*’*). (D-D*’*)ppr*^*A*^ mutant clones (non green cells, *D* and dashed white line, *D*’) on overexpression of *Prosβ6*^*1*^ under *Actin*>*Gal4*, wing discs immunostained for Marf::HA (red). *(E)*Quantification for relative fluorescence intensities of Marf::HA in *ppr*^*A*^ mutant clones on treatment with DMSO (n=24), chloroquine (n=6), MG132 (n=10) and on overexpression of *Prosβ6*^*1*^ under *Actin*>*Gal4* (n=15). Graphs represent average intensity values normalized to that of control cells. Two tailed paired t-test between control and *ppr*^*A*^ mutant cells. Significance represented by p<0.001***

To investigate the role of UPS in Marf downregulation in *ppr* mutants, we fed larvae with the proteasomal inhibitor MG132 (40,41). While in DMSO-fed larvae, *ppr*^*A*^ mutant clones had lower levels of Marf:HA as compared to the neighboring wild type cells, MG132-fed larvae show no change in Marf::HA levels (Figure 2A-A’, 2C-C’, 2E). We further expressed a dominant negative form of *Pros 6* to inhibit UPS activity (42) and tested its effect on Marf::HA levels in *ppr*^*A*^ mutant clones. Similar to MG132 treatment, we found that Marf::HA levels were restored in *ppr*^*A*^ mutant clones upon *Pros 6*^*1*^ overexpression (Figure 2D-D’, 2E). These results suggest that UPS-mediated degradation of Marf results in Marf downregulation in *ppr*^*A*^ mutant clones.

### PINK1 and Park dependent *Marf* regulation in *ppr* mutants

Several E3 ubiquitin ligases have been linked to Mitofusin degradation. For example, Mitofusin degradation by HUWE1 occurs under genotoxic stress or on altered fat metabolism (43,44) while Mitofusin degradation by Park occurs upon mitochondrial membrane depolarization (45,46). In *Drosophila* too, HUWE1, MUL1 and Park have been shown to affect Marf levels (27,43,47). Similar to our observation in *ppr*^*A*^ mutant clones (Fig. 3A-A’, 3D), we found a down regulation of Marf::HA levels in *ppr*^*A*^ *HUWE1*^*B*^ double mutant clones (Figure S2A-A’) and *ppr*^*A*^ mutant clones in *MUL1*^*A6*^ mutant background (Figure S2B-B’). Interestingly, *ppr*^*A*^ mutant clones in *park*^*Δ21*^ mutant background did not show Marf::HA down regulation suggesting Park, but not HUWE1 and MUL1, is required for Marf down regulation in *ppr* (Figure 3B-B’, 3D). Since *park* is genetically downstream to *Pink1 (48,49)*, we tested whether PINK1 is also required for Marf degradation in *ppr* mutant clones. We generated *ppr*^*A*^ *Pink1*^*5*^ double mutant clones and found that these clones do not show a reduction in Marf::HA levels (Figure 3C-C’, 3D), suggesting that mitochondrial impairment in *ppr* mutant cells causes PINK1-Park activation, and subsequently, Marf downregulation. Our observations match previous reports of down regulation of Mfn1 and Mfn2 upon CCCP treatment as a mechanism to suppress mitochondrial fusion prior to PINK1-Park mediated mitophagy (45,46).

**Figure 3.**
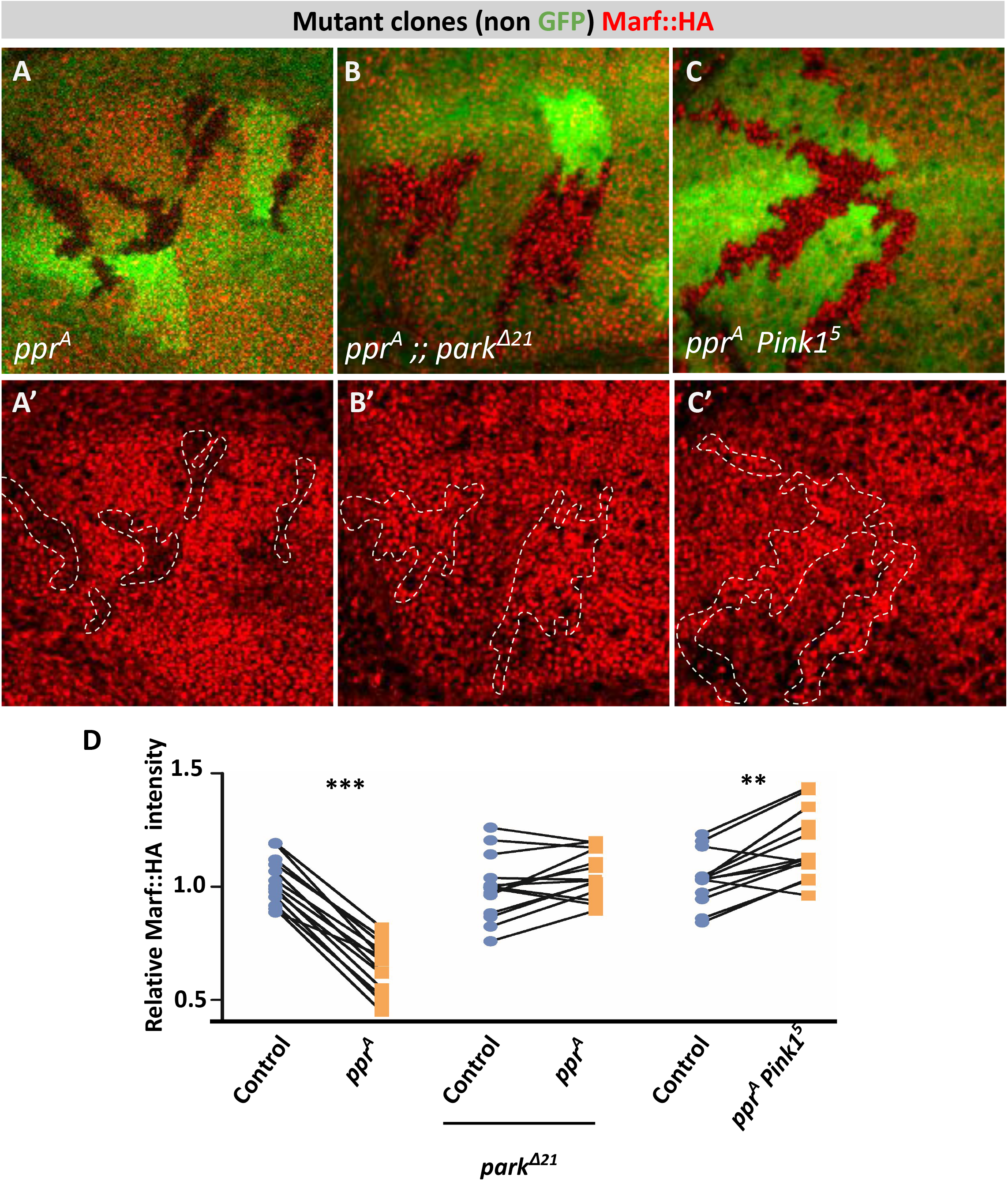
PINK1-Park are required for Marf degradation in *ppr* mutants: *(A-A*’*)ppr*^*A*^ mutant clones (non green cells, *A* and dashed white line, *A*’), wing discs immunostained for Marf::HA (red). *(B-B*’*)ppr*^*A*^ mutant clones (non green cells, *B* and dashed white line, *B*’) in *park*^*Δ21*^ background, wing discs immunostained for Marf::HA (red). *(C-C*’*)ppr*^*A*^ *Pink1*^*5*^ double mutant clones (non green cells, *C* and dashed white line, *C*’), wing discs immunostained for Marf::HA (red). *(D)*Quantification for relative fluorescence intensities of Marf::HA in *ppr*^*A*^ mutant clones (n=14), *ppr*^*A*^ mutant clones in *park*^*Δ21*^ background (n=17) and *ppr*^*A*^ *Pink1*^*5*^ double mutant clones (n=13). Graphs represent average intensity values normalized to that of control cells. Two-tailed paired t-test between control and mutant cells. Significance represented by p<0.01**, p<0.001***

### UPR^my^ is not sufficient to induce Marf downregulation in *ppr* mutants

The role of the PINK1-Park pathway in mitochondrial quality control is well known. However, the exact mechanism of PINK1-Park activation in *in vivo* contexts remains unclear. In cancer cell lines, dissipation of MMP and increased oxidative stress have been shown to activate PINK1-Park on the OMM leading to mitophagy (10,50). Therefore, oxidative stress or reduced MMP may activate PINK1-Park and subsequent Marf degradation in *ppr* mutants. However, we have shown earlier that *ppr* mutants do not have increased oxidative stress as compared to controls (29). We checked MMP in *ppr*^*A*^ mutant clones using TMRE, a dye that reversibly stains mitochondria in a membrane potential-dependent manner. We observed that TMRE intensity in *ppr*^*A*^ mutant clones is similar to that of wild type cells (Figure S3A-A’). These observations rule out the possibility that PINK1-Park is activated due to oxidative stress or altered MMP in *ppr* mutants.

Mitochondrial unfolded protein response (UPR^mt^), which is a cellular response to altered mitochondrial proteostasis, has been shown to activate PINK1-Park leading to mitophagy (51). Therefore, we first tested for UPR^mt^ activation in *ppr* mutants. We determined the levels of Hsp60A which is reported to be increased due to elevated UPRmt (52). We found increased Hsp60A levels in ppr mutants as compared to controls, suggesting elevated UPRmt in ppr mutants (Figure S3B-B’). Activation of UPR^mt^ upon the loss of *LRPPRC* has also been observed in *C*.*elegans* and mammalian cells, and hence, it appears to be an evolutionarily conserved phenomenon (53). Increased UPR^mt^ may induce PINK1-Park activity, which in-turn could lead to Marf downregulation. Therefore, we genetically suppressed the UPR^mt^ response pathways and checked its impact on Marf in *ppr* mutants. Transcription factors Crc (homolog of ATF4), Dve and Foxo are known to mediate UPR^mt^ (54–56). We generated *ppr*^*A*^ mutant clones with either *crc, foxo* or *dve* knocked down. None of these interventions affected Marf::HA downregulation in *ppr*^*A*^ clones, suggesting that the activation of these UPR^mt^ pathways may not be causing PINK1-Park activation (Figure S3C-E’). However, these interventions would not change the altered mitochondrial proteostasis in *ppr* mutants, which can activate PINK1-Park. Since, to the best of our knowledge, there is no reported method to suppress mitochondrial proteostasis defects, we asked whether the induction of mitochondrial proteostasis defects is sufficient to induce Marf degradation. To induce mitochondrial proteostasis defects, we expressed a mutant form of ornithine transcarbamylase (ΔOTC) that accumulates in an unfolded state and is shown to trigger UPR^mt^ in flies (56). We expressed either ΔOTC or wild type OTC in the posterior half of the wing disc using *En*-*Gal4* (*En*>*Gal4*/+; *UAS-ΔOTC*/+ or *En*>*Gal4*/+; *UAS-OTC*/+) and tested Marf::HA levels. We found neither OTC nor ΔOTC expression changed Marf::HA levels in the posterior half (marked by RFP) as compared to the anterior half of the wing discs (Figure S3F-F’, S3G-G’). Although these observations do not rule out a role for mitochondrial proteostasis in activating PINK1-Park in *ppr* mutants, our data suggest that UPR^mt^ is not sufficient to cause Marf degradation.

### Bendless, a K63-linked E2 ubiquitin conjugase, is a regulator of Marf

To gain further insight into PINK1-Park activation and Marf degradation, we screened for a gene whose loss may cause a subtle increase in Marf levels as observed in *park*^*Δ21*^ and *Pink1*^*5*^ mutant clones (Figure S4A-A’, S4B-B’). In our genetic screen, we found two independent alleles of *bendless* (*ben*^*A*^ and *ben*^*B*^) showing a subtle but consistent increase in Marf::HA levels in mutant clones (Figure 4A-A’, 4E, S4C-C’). This was also confirmed by western blot using whole larval extracts (Figure 4G-G”). Ben is a fly homologue of the K63-linked E2 ubiquitin conjugase UBE2N/UBC13 with a marked similarity from yeast to humans (Figure S4G). We ruled out the possibility that the increase in Marf::HA levels upon the loss of Ben is due to increased mitochondrial content by determining Tom20 levels, as there was no difference in Tom20::mCherry levels between *ben* mutant clones and controls (Figure 4B-B’, 4E). We also did not find an increase in *Marf* mRNA levels in *ben* mutants suggesting that the increase in Marf protein levels is not a consequence of increased transcription (Figure 4F). These data suggest that Ben regulates Marf levels post-transcriptionally.

**Figure 4:**
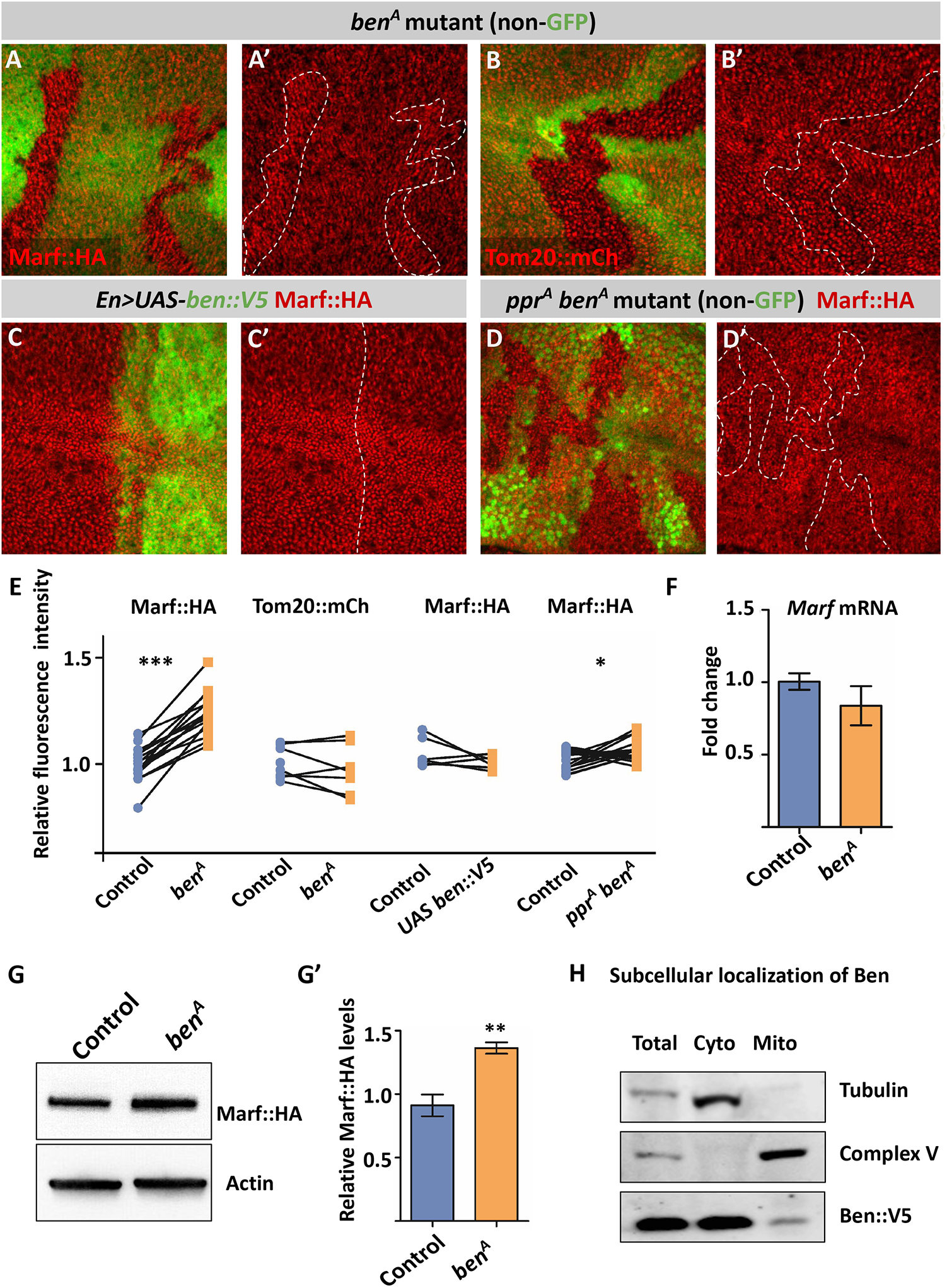
Ben is required for Marf degradation in *ppr* mutants: *(A-B*’*)ben*^*A*^ mutant clones (non green cells, *A, B* and dashed white line, *A*’, *B*’), wing discs immunostained for Marf::HA (red, *A-A*’), Tom20::mCh (red, *B-B*’). *(C-C*’)Overexpression of *ben::V5* using *En*>Gal4, wing discs immunostained for Ben::V5 (green) and Marf::HA (red). *(D-D*’*)ppr*^*A*^ *ben*^*A*^ double mutant clones (non green cells, *D* and dashed white line, *D*’), wing discs immunostained for Marf::HA (red). *(E)*Quantification for relative fluorescence intensities of Marf::HA in *ben*^*A*^ mutant clones (n=15), Tom20::mCh in *ben*^*A*^ mutant clones (n=7), Marf::HA on overexpression of *ben::V5* (n=6), Marf::HA in *ppr*^*A*^ *ben*^*A*^ double mutant clones (n=16). Graphs represent average intensity values normalized to that of control cells. Two-tailed paired t-test between control and mutant cells. *(F)*Quantification of *Marf* mRNA levels in third instar *ben* mutant (*y w ben*^*A*^ FRT19A) larvae compared to control *(y w* FRT19A) (n=3). Two tailed unpaired t-test between control and *ben*^*A*^ mutant larvae. *(G-G*’)Representative western blot for *ben* mutant (*y w ben*^*A*^ FRT19A) and control (*y w* FRT19A) larval lysate probed for Marf::HA and Actin. *(G*’)Quantification for intensity of Marf::HA band normalized to Actin band intensity for *ben*^*A*^ mutant and control larvae (n=5). Two-tailed unpaired t-test between control and *ben*^*A*^ mutant larvae. *(H)*Representative western blot for total larval lysate (Total), Cytoplasmic fraction (Cyto), and Mitochondrial enriched fraction (Mito) probed for Ben::V5, Complex V and Tubulin. Ben::V5 is present in Total and Cyto fraction in abundance, while a small amount of Ben::V5 is visible in Mito fraction. Error bars represent S.E.M. Significance represented by p<0.05*, p<0.01**, p<0.0001***

Next, we asked whether Ben is sufficient to induce Marf degradation. To test this, we generated a C-terminal V5-tagged Ben *(UAS-ben::V5)* transgenic line for tissue specific expression of *ben* and confirmed that the fusion protein is biologically functional by complementing the lethality associated with the *ben*^*A*^ mutant allele (Figure S4F, S4H). We then expressed *ben::V5* in the posterior half of the wing disc using the *En*-*Gal4* driver and compared the fluorescence intensities of Marf::HA in the posterior and the anterior halves. We found that *ben::V5* overexpression did not affect the levels of Marf::HA (Figure 4C-C’, 4E). Additionally, we overexpressed an N-terminal HA-tagged Ben (*UAS-HA::ben*) using *En*-*Gal4* and found no change in Marf::mCherry levels (Figure S4E-E’). These data suggest that Ben is necessary but not sufficient for Marf downregulation. Since loss of *ben, Pink1* or *park* results in mild upregulation of Marf, we hypothesize that Ben acts in the PINK1-Park pathway to regulate the steady state levels of Marf.

### Bendless is essential for Marf downregulation in *ppr* mutants

Given that Marf undergoes proteolytic degradation in *ppr* mutants, we wanted to look at if Ben is involved in Marf degradation not only basally but under mitochondrial stress as well. We thus created *ppr* and *ben* double mutant clones and found that *ppr*^*A*^ *ben*^*A*^ or *ppr*^*A*^ *ben*^*B*^ double mutant clones showed no reduction in Marf::HA levels, unlike *ppr* mutant clones (Figure 4D-D’, 4E and S4D-D’). This suggests that Ben is essential for Marf degradation in *ppr* mutant cells.

### Bendless is required for PINK1 stability and activity

To understand the mechanism of Ben mediated Marf degradation, we first looked at subcellular localization of Ben using the Ben::V5 tagged line. We overexpressed Ben::V5 using *Act*-*Gal4*, and performed cell fractionation from whole larval extracts. We found that in addition to the cytoplasmic fraction (marked by the presence of Tubulin) Ben::V5 is also present in the mitochondria enriched fraction (marked by presence of ComplexV) (Figure 4H). Mitochondria Ben might regulate the activity PINK1-Park pathway to degrade Marf. To study the role of Ben in the PINK1-Park pathway, we tested the functional interaction between *ben* and *Pink1* with respect to Marf degradation. Since PINK1-Park activity is suppressed by PINK1 degradation (57), PINK1 overexpression may activate PINK1-Park mediated Marf downregulation in the wing disc. We overexpressed *Pink1* in the posterior half of the discs (*En*-*Gal4*/*UAS*-*Pink1, UAS-RFP* or *UAS-GFP*) and checked the levels of Tom20::mCherry, Complex-V and Marf::HA. We observed no change in Tom20::mCherry and Complex-V levels (Figure 5A-A’, 5D, S5A-A’), but a marked reduction in Marf::HA levels (Figure 5B-B’), suggesting that PINK1 is both necessary and sufficient to downregulate Marf. *Pink1* overexpression in the wing discs may not have a significant impact on mitophagy and thus the mitochondrial content (Tom20::mCherry and Complex-V) is not affected. To test the functional interaction between Ben and PINK1, we created *ben*^*A*^ mutant clones in both wild type and in *Pink1* overexpression backgrounds and found that *Pink1* overexpression does not induce Marf::HA downregulation in *ben*^*A*^ mutant clones (Figure 5C-C’, 5D) or in *ben*^*A*^ mutant wing discs (Figure S5B-B’). These data suggest that Ben is necessary for PINK1 activity to cause Marf downregulation.

**Figure 5.**
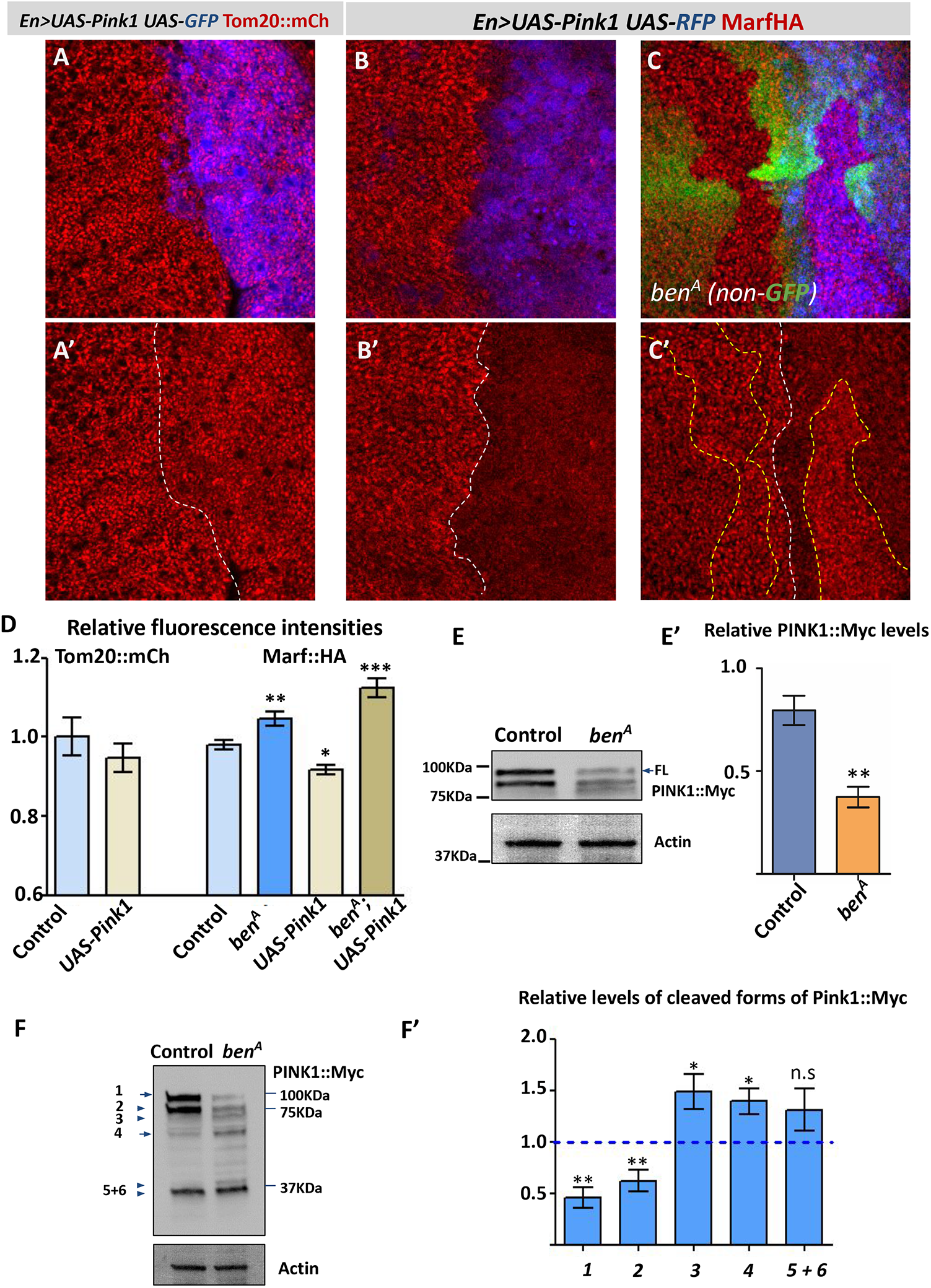
*ben* is required for PINK1 mediated Marf degradation: *(A-A*’)Overexpression of *Pink1* using *En*>*Gal4*, wing discs marked with *UAS-RFP* (blue, *A*) and immunostained for Tom20::mCh (red, *A-A*’). *(B-B*’)Overexpression of *Pink1* using *En*>*Gal4*, wing discs marked with *UAS-RFP* (blue, *B*) and immunostained for Marf::HA (red, *B-B*’). *(C-C*’*)ben*^*A*^ mutant clones (non green cells, *C* and dashed yellow line, *C*’) in background of overexpression of *Pink1* using *En*>*Gal4*, wing discs marked with *UAS-RFP* (blue, *C*) and immunostained for Marf::HA (red, *B-B*’). *(D)*Quantification for relative fluorescence intensities of Tom20::mCh in *UAS-Pink1* cells (n=7), Marf::HA in *ben*^*A*^ mutant clones (n=17), *UAS-Pink1* cells (n=8) and *ben*^*A*^ mutant clones in *UAS-Pink1* background (n=8). Graphs represent average intensity values normalized to that of control cells. Two-tailed paired t-test between control and mutant cells or cells Overexpression of *Pink1. (E-E*’)Representative western blot for *ben* mutant (*y w ben*^*A*^ FRT19A) and control (*y w* FRT19A) larval lysate probed for Myc and Actin. *(E*’)Quantification for intensity of full length (FL) PINK1::Myc band normalized to actin band intensity for *ben*^*A*^ mutant and control larvae (n=8). Two-tailed unpaired t-test between control and mutant larvae. *(F-F*’)Representative western blots for *ben* mutant (*y w ben*^*A*^ FRT19A) and control (*y w* FRT19A) larval lysate probed for Myc and Actin. *(F*’)Quantification for intensity of cleaved PINK1::Myc bands normalized to actin band intensity for *ben*^*A*^ mutant and control larvae (n=8). One sample two-tailed t-test of bands of *ben*^*A*^ mutant larvae normalized to its corresponding control band taken as one. Same samples were used to quantify results as in Fig 5E’ and Fig 5F’. Error bars represent S.E.M. Significance represented by p<0.05*, p<0.01**, p<0.0001***, n.s - non significant.

To understand how Ben may regulate PINK1 activity, we checked the effect of loss of Ben on PINK1 levels. We performed western blots using whole larval extracts from control and *ben*^*A*^ mutants containing genomic tagged PINK1::Myc. We found a significant downregulation of full length PINK1::Myc in *ben*^*A*^ mutants, but an increase in low molecular weight PINK1::Myc bands, suggesting that Ben is required for stability of full length PINK1 (Figure 5E-F’). The low molecular weight bands might be products of PINK1 degradation by mitochondrial proteases as described by Thomas et.al. (58). Taken together, our data suggests that Ben is required for the stability of PINK1 and mediates the homeostatic turnover of Marf.

### *Ben* regulates mitochondrial dynamics under mitochondrial stress

An RNAi-based screen in larval fat bodies has shown *ben* RNAi leads to enlarged mitochondria, similar to when *Pink1* or *park* are knocked down (59). This would be in accordance with our observation wherein loss of ben results in Marf upregulation. To better characterize mitochondrial morphology we looked at Complex-V antibody staining in mutant larval muscles. In this tissue, loss of ben shows a similar filamentous network as seen in wildtype (Figure 6A-B, S6A-B). However, loss of *ben* in *ppr* mutants exacerbates mitochondrial morphology defects seen in ***ppr*. On loss of *ppr* alone larval muscles show distinctive large globular mitochondria along with filamentous and ring-shaped mitochondria (Figure 6C**, S6C). *ppr ben* double mutants rarely show filamentous mitochondria (Figure 6D). Instead, we observed a significant increase in the size and frequency of large globular and ring-shaped mitochondria as compared to *ppr* (Figure 6D). We also observed that in lesser frequency *ppr*^*A*^ *ben*^*A*^ double mutants mitochondria form clusters, especially around the nucleus which is not observed in either *ppr*^*A*^ or *ben*^*A*^ mutants (Figure 6D, S6D). These results suggest that Ben is required to suppress the hyperfusion of defective mitochondria in *ppr* and thereby regulates mitochondrial quality control.

**Figure 6.**
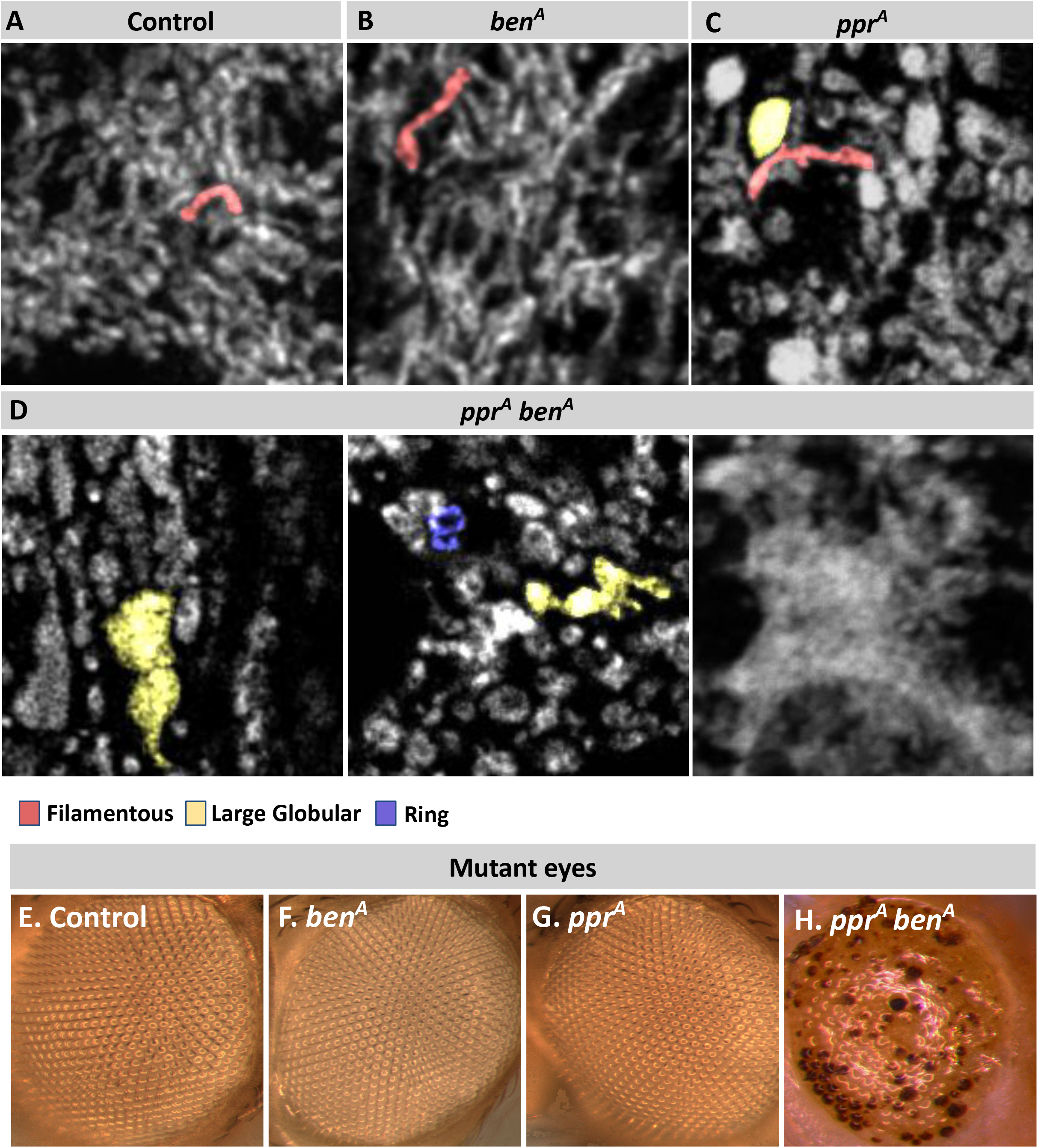
Ben is required for maintaining mitochondrial morphology and neuronal health in *ppr* mutants: *(A-D)*Confocal sections of 3^rd^ instar larval muscles immunostained for Complex-V (gray) in control*(A), ben*^*A*^*(B), ppr*^*A*^*(C)* and *ppr*^*A*^ *ben*^*A*^*(D)* larvae. Representative individual mitochondrial morphology is marked by different colors: filamentous (red), large globular (yellow) and ring (blue).*(E-H)*Mutant eye clones from young flies of control*(E), ben*^*A*^*(F), ppr*^*A*^*(G)*, and *ppr*^*A*^ *ben*^*A*^*(H)* genotypes.

### Loss of Bendless accelerates photoreceptor degeneration in *ppr* mutants

Mutations in human *LRPPRC* cause Leigh Syndrome, a neurometabolic disease (60). Earlier work has shown that mutations in *ppr* cause activity induced retinal degeneration (29). As mutations in *ben* exacerbates the mitochondrial morphology phenotypes in *ppr* mutants, loss of *ben* may accelerate the degenerative phenotype. To test this hypothesis, we made eye specific *ppr, ben* and *ppr ben* double mutant clones using the *ey-FLP* system (30). We found that *ppr* mutant and *ben* mutant eyes show normal morphology but *ppr ben* double mutant eyes show severe retinal degeneration in young flies suggesting that loss of *ben* can accelerate retinal degeneration (Figure 6E-H). This result suggests that Marf regulation by Ben is a neuroprotective mechanism.

## Discussion

To identify novel regulators of mitochondrial fusion in an *in vivo* system, we screened fly mutants for altered Marf levels. We found that mutations in *ppr*, which result in mitochondrial dysfunction, causes reduction in Marf levels (Figure 1A). We found that in *ppr* mutants, Marf is degraded by the UPS in a PINK1-Park dependent mechanism (Figure 2C-D, 3A-C). In the screen, we also identified mutations in *ben* causing subtle Marf upregulation (Figure 4A, 4G). We found that Ben is essential for PINK1 activity (Figure 5C), regulates Marf levels (Figure 4D) and mitochondrial morphology (Figure 6D) in *ppr* mutants. We also found that a loss-of-function mutation of both *ppr* and *ben* in the eyes results in accelerated retinal degeneration (Figure 6H) indicating that under mitochondrial stress Ben mediated regulation of mitochondrial dynamics is a protective mechanism (Figure 7B).

**Figure 7:**
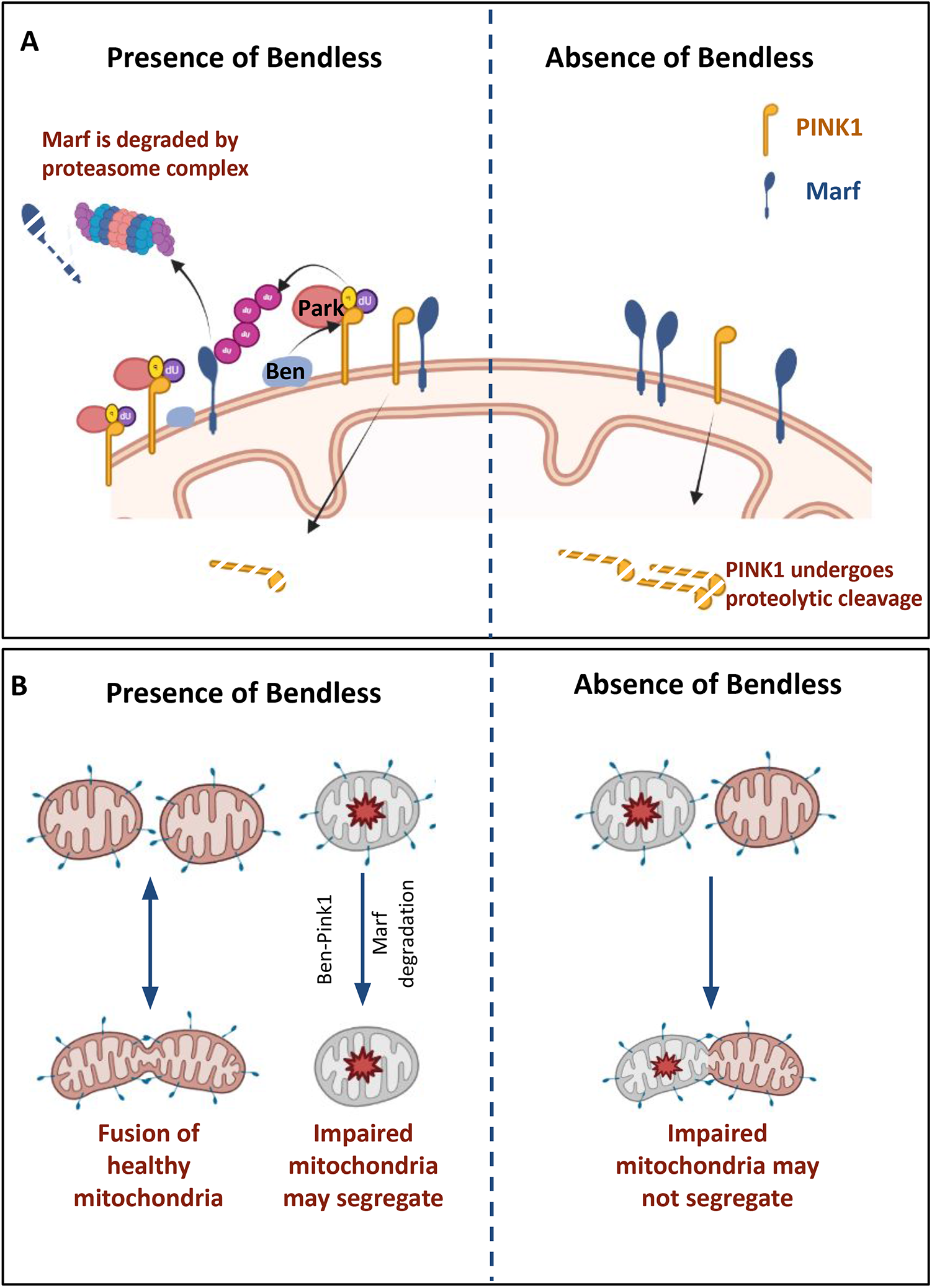
*(A)***Mitofusin turnover is regulated by Bendless mediated stabilization of PINK1.** In presence of Ben, PINK1 is stabilized on the mitochondrial surface and results in Park mediated ubiquitination of Marf.Ubiquitinated Marf is degraded by the proteasome complex. Absence of Ben results in increased proteolytic cleavage of PINK1 resulting in accumulation of Marf. *(B)***Bendless prevents fusion of impared mitochondria**. In healthy mitochondria Ben-PINK1 mediated Marf turnover keeps adequate Marf protein aiding mitochondrial fusion. Upon mitochondrial stress (as we show in case of mutation in *ppr*) results in presence of impaired mitochondria, we predict that the increase in Ben-PINK1 mediated Marf degradation keeps the damaged mitochondria isolated and thus can be repaired or sent for mitophagy. In the absence of Ben however Marf is not removed from impaired mitochondria and results in fusion of impaired mitochondria with the healthy pool.

Increased mitochondrial size has been observed in several mitochondrial diseases (61); however, it is not clear as to how these abnormal mitochondria contribute to disease progression. Mitochondrial fusion occurs in a bid to increase oxidative phosphorylation under various cellular and mitochondrial stresses, a response termed as stress induced mitochondrial hyperfusion [SIMH (3)] (62,63). We propose that reduced ETC activity and mitochondrial stress in *ppr* (28,29,36), can induce SIMH (Figure 6D, S1E). Mitochondrial fusion has been observed upon loss of *ppr* homologs in *C*.*elegans*, mouse and human cell lines (35,64) as well as in other mutants where the ETC is compromised (52,65,66). Since SIMH increases ATP synthesis and inhibits mitophagy (3,4,67,68), increased mitochondrial size appears to be a compensatory adaptation in *ppr* mutants in response to a bioenergetic deficit or mitochondrial stress.

Despite the increased mitochondrial size in *ppr* mutants (Figure S1E), we observed Marf downregulation. We hypothesize that, while an adaptive mechanism may induce SIMH *(cellular response)*, MQC may induce Marf degradation to suppress fusion of dysfunctional mitochondria (*mitochondrial response*) (Figure 7B). As proposed earlier (62,63), this condition appears to be a conflict between the bioenergetic adaptations and the MQC mechanisms. Alternatively, increased mitochondrial size in *ppr* mutant cells may induce PINK1-Park to suppress further mitochondrial fusion by Marf degradation. A similar hypothesis was also proposed by Yamada et al. (69). We found that Marf degradation in *ppr* mutant clones in developing wing primordium is dependent on Park (Figure 3B). We also observed a subtle increase in Marf levels in *park* and *Pink1* mutants (Figure S4A-B) (27). This suggests that the PINK1-Park plays a homeostatic role in Marf turnover in wild type tissue, while mitochondrial impairments —as in *ppr* mutants (29)— may further amplify its activity to reduce Marf levels (Figure 1A) possibly to segregate damaged mitochondria (9,45,46). The remarkable discovery by Narendra et al. that CCCP, which dissipates MMP, induces PINK1-Park-dependent mitophagy in cancer cells provided an unparalleled assay to investigate the mechanism further (10,70). However, we observe PINK1-Park activation in the absence of severe mitochondrial depolarisation (Figure S3A) in *ppr* mutants results in Marf degradation and not mitophagy (Figure 1A-B). We also find the PINK1 overexpression is sufficient to induce Marf degradation without triggering mitophagy (Figure 5A-B, S5A). *In vivo* studies have shown PINK1-Park to function both in mitophagy (13–20) and mitochondrial dynamics (22–26), but the physiological or cellular contexts that may determine various downstream activities of PINK1-Park are not known (21,71,72). Thus *ppr* mutants provide a novel and physiologically relevant *in vivo* system to study PINK1-Park mediated Marf regulation under mitochondrial stress.

In steady state conditions, PINK1 is imported into the mitochondria and cleaved by mitochondrial peptidases, it then retro translocates to the cytoplasm and is degraded by UPS to limit PINK1-Park activity (58,73,74). Loss of MMP, increased oxidative stress or increased UPR^mt^ stabilizes full length PINK1, which then recruits Park leading to ubiquitination of OMM proteins and mitophagy (9,10,51,75,76). Given no change in MMP (Figure S3A) and oxidative stress in *ppr* mutants (28,29,37), we suspected that mitochondrial proteostasis activates PINK1-Park to downregulate Marf. However, activation of UPR^mt^ by ΔOTC expression did not result in Marf degradation suggesting that activation of UPR^mt^ alone may not be sufficient to activate PINK1-Park mediated Marf degradation *in vivo* (Figure S3G). Identification of additional factors leading to PINK1-Park activation for Marf degradation *in vivo* requires further investigation.

In most cells, PINK1 activity is maintained at low levels via its constant turnover by mitochondrial proteases and the UPS (57,77). For example, CHIP-mediated K48-ubiquitination promotes PINK1 turnover (78), while BAG2, a chaperon, prevents ubiquitination and promotes PINK1 stability (79,80). We found that Marf degradation in *ppr* mutants or by *Pink1* overexpression is completely suppressed in the absence of the K63-linked E2 conjugase Ben (Figure 4D, S4D and 5C). Previous studies have observed that the mammalian homolog of Ben, UBE2N, is dispensable for mitophagy but facilitates the clustering of mitochondria during CCCP induced mitophagy (81–83). We also found that the loss of *ben* does not alter developmental mitophagy during larval midgut remodeling (Figure S5C), which has been shown to be dependent on PINK1-Park **(13,14)**.

We hypothesize three possible mechanisms through which Ben regulates Marf degradation. One, given that Park activity is regulated by K63 ubiquitination (84), Ben may ubiquitinate Park. Two, Ubiquitin C-terminal hydrolase L1 (UCHL1), which suppresses Marf degradation (85), is K63 ubiquitinated leading to its autophagic degradation (86). Thus, Ben might mediate K63 ubiquitination and degradation of UCHL1. K63 ubiquitination of PINK1 by the Traf6-SARM1 complex is shown to stabilize PINK1 (87). Therefore, Ben may stabilize PINK1 by K63 ubiquitination. The fact that loss of *ben* results in reduced PINK1 levels (Figure 5E-E’), suggests Ben is likely to increase the stability of PINK1 by K63 ubiquitination. Indeed human PINK1, in cell culture systems, is known to be ubiquitinated at K137 by both K48 and K63 linkages (88). While K48 chains are linked with PINK1 degradation; the significance of K63 linkage is not obvious. K63 ubiquitination is suggested to protect proteins from proteasomal degradation (89). Overall, Ben-mediated K63 ubiquitination appears to be responsible for PINK1 stability (Fig. 7A).

In conclusion, Ben-PINK1-Park regulation of Marf appears to be a homeostatic function which is further activated in response to aberrant mitochondrial function. *ppr ben* double mutants show aberrant mitochondrial morphology in larval muscles and severe retinal degeneration as compared to *ppr* mutant eyes indicating a protective role for Ben (Figure 6). Given that mutations in *LRPPRC* result in Leigh syndrome, it would be crucial to check the activation of Ben-PINK1-Park in Leigh syndrome and other mitochondrial diseases. Indeed, altered mitochondrial dynamics has been reported in many mitochondrial diseases (61,90–92). It is possible an adaptive response in these diseases can modify mitochondrial dynamics in a Ben-PINK1-Park-dependent mechanism. Thus, further studies on the mechanisms of Ben-PINK1-Park activation will be crucial for understanding mitochondrial quality control in mitochondrial disease.

## Figure Legends

**Supplementary Figure 1:**
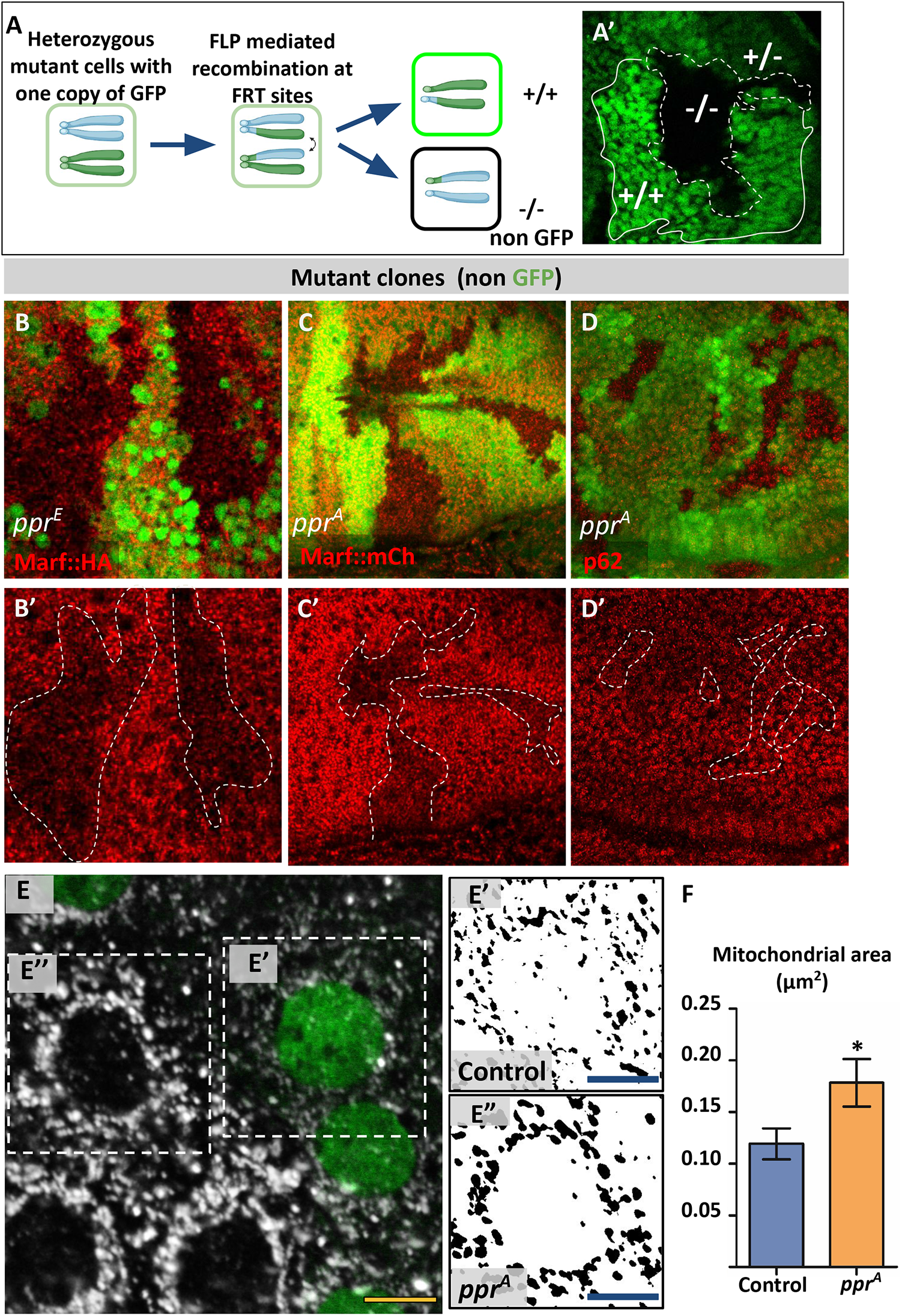
(*A)*Schematic to illustrate Flp-FRT mediated recombination system. *(A*’) Green marks wild type and heterozygous cells (solid white line, +/+ and +/-) absence of GFP marks mutant clones/cells (dashed white line, -/-). *(B-B*’*)ppr*^*E*^ mutant clones (non green cells, *B* and dashed white line, *B*’), wing discs immunostained for Marf::HA (red). (*C-C*’*)ppr*^*A*^ mutant clones (non green cells, *C* and dashed white line, *C*’), wing discs immunostained for Marf::mCh (red). *(D-D*’*)ppr*^*A*^ mutant clones (non green cells, *D* and dashed white line, *D*’), wing discs immunostained for endogenous p62 (red). *(E-E*’’*)ppr*^*A*^ mutant clones (non green cells, *E*) in peripodial cells of third instar larval wing discs, immunostained for Complex-V (gray). Inset of control *(E*’) and *ppr*^*A*^ mutant cell *(E*”) from E. (E’-E’’). Binary image of Complex-V staining. Scale bar represents 10µm in *(A)* and 4µm in *(E*’ and *E*”).

**Supplementary Figure 2:**
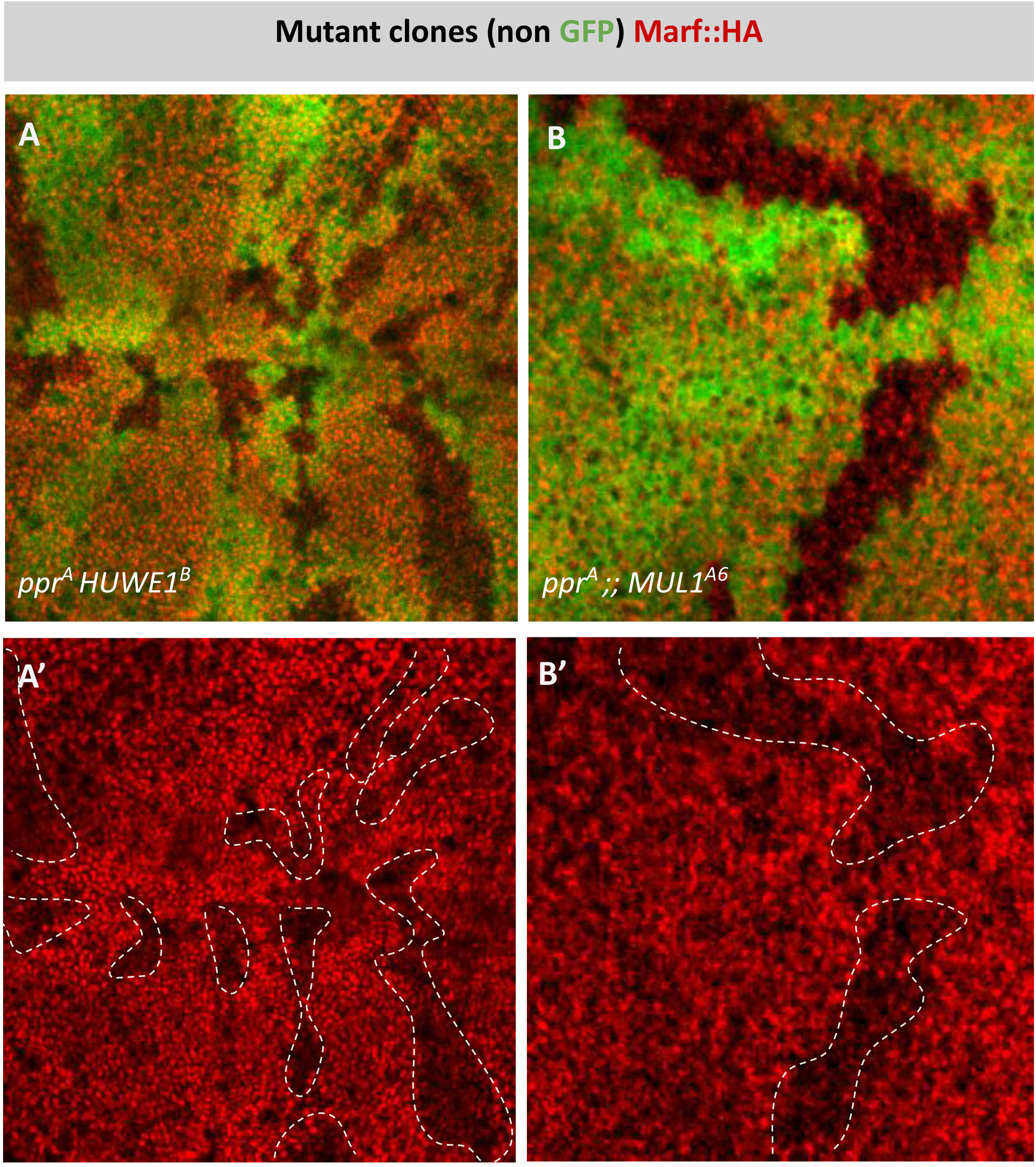
*(A-A*’*)ppr*^*A*^ *HUWE1*^*B*^ double mutant clones (non green cells, *A* and dashed white line, *A*’), wing discs immunostained for Marf::HA (red). *(B-B*’*)ppr*^*A*^ mutant clones (non green cells, *B* and dashed white line, *B*’) in *MUL1*^*A6*^ mutant background, wing discs immunostained for Marf::HA (red).

**Supplementary Figure 3:**
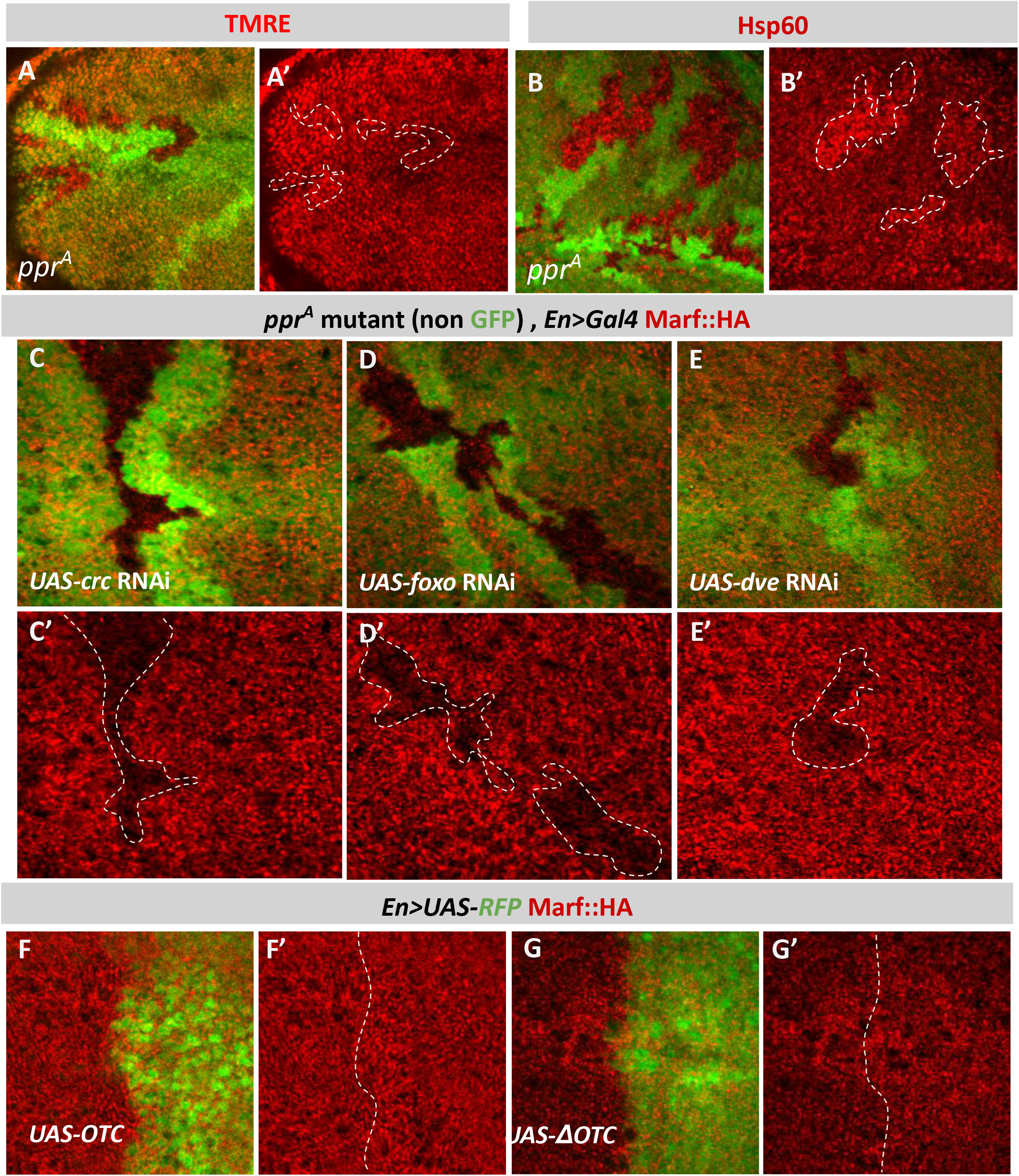
*(A-A*’*)ppr*^*A*^ mutant clones (non green cells, *A* and dashed white line, *A*’), wing discs stained for TMRE (red) and live imaged. *(A*”)Quantification for relative fluorescence intensity of TMRE in *ppr*^*A*^ mutant clones (n=20). Graphs represent average intensity values normalized to that of control cells. Two-tailed paired t-test between control and *ppr*^*A*^ mutant cells showed no significant change. *(B-B*’*)ppr*^*A*^ mutant clones (non green cells, *B* and dashed white line, *B*’), wing discs immunostained for Hsp60 (red).*(C-D*’*)ppr*^*A*^ mutant clones (non green cells, *C, D, E* and dashed white line, *C*’ *D*’, *E*’) on KD of *crc(C-C*’*), foxo(D-D*’) and *dve(E-E*’) using *En*>Gal4, wing discs marked by *UAS-RFP* (green) and immunostained for Marf::HA (red).*(F-G*’)Overexpression of *OTC(F-F*’) and *ΔOTC(G-G*’) using *En*>Gal4, wing discs marked by *UAS-RFP* (green) and immunostained for Marf::HA (red).

**Supplementary Figure 4:**
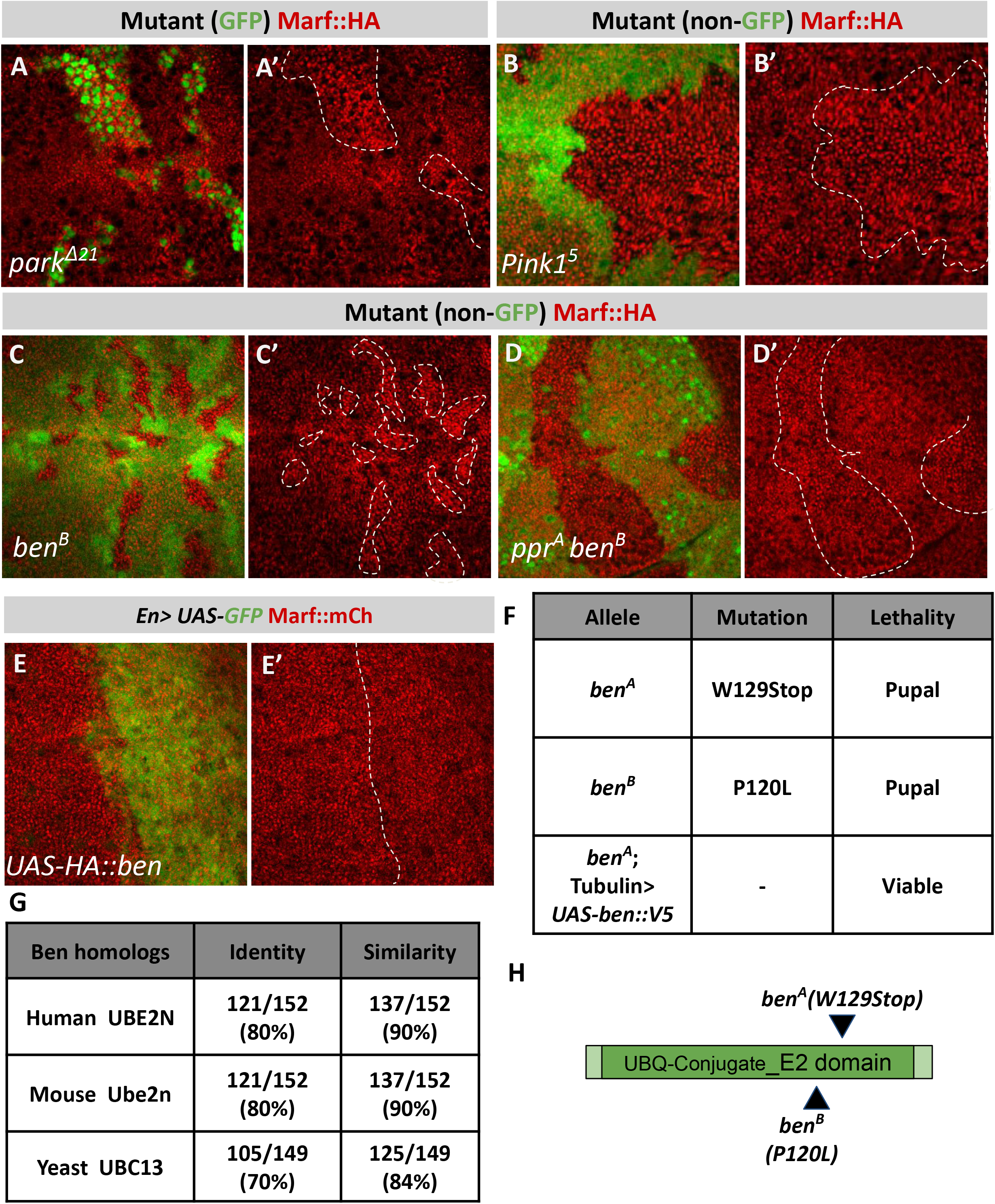
*(A-E*’)Wing discs immunostained for Marf::HA (red) in *park*^*Δ21*^ mutant clones (green cells, *A-A*’*) Pink1*^*5*^ mutant clones (non green cells, *B-B*’*), ben*^*B*^ mutant clones (non green cells, *C-C*’) and *ppr*^*A*^ *ben*^*B*^ (non green cells, *D-D*’). Wing discs immunostained for Marf::mCh (red) on overexpression of HA::Ben using *En*>Gal4 wing discs marked with *UAS-GFP* (green) *(E-E*’*). (F)*Table for Ben mutations and lethal staging. *(G)*Similarity and identity between Ben and its homologs. *(H)*Schematic showing point mutations in *ben*^*A*^ and *ben*^*B*^ alleles.

**Supplementary Figure 5:**
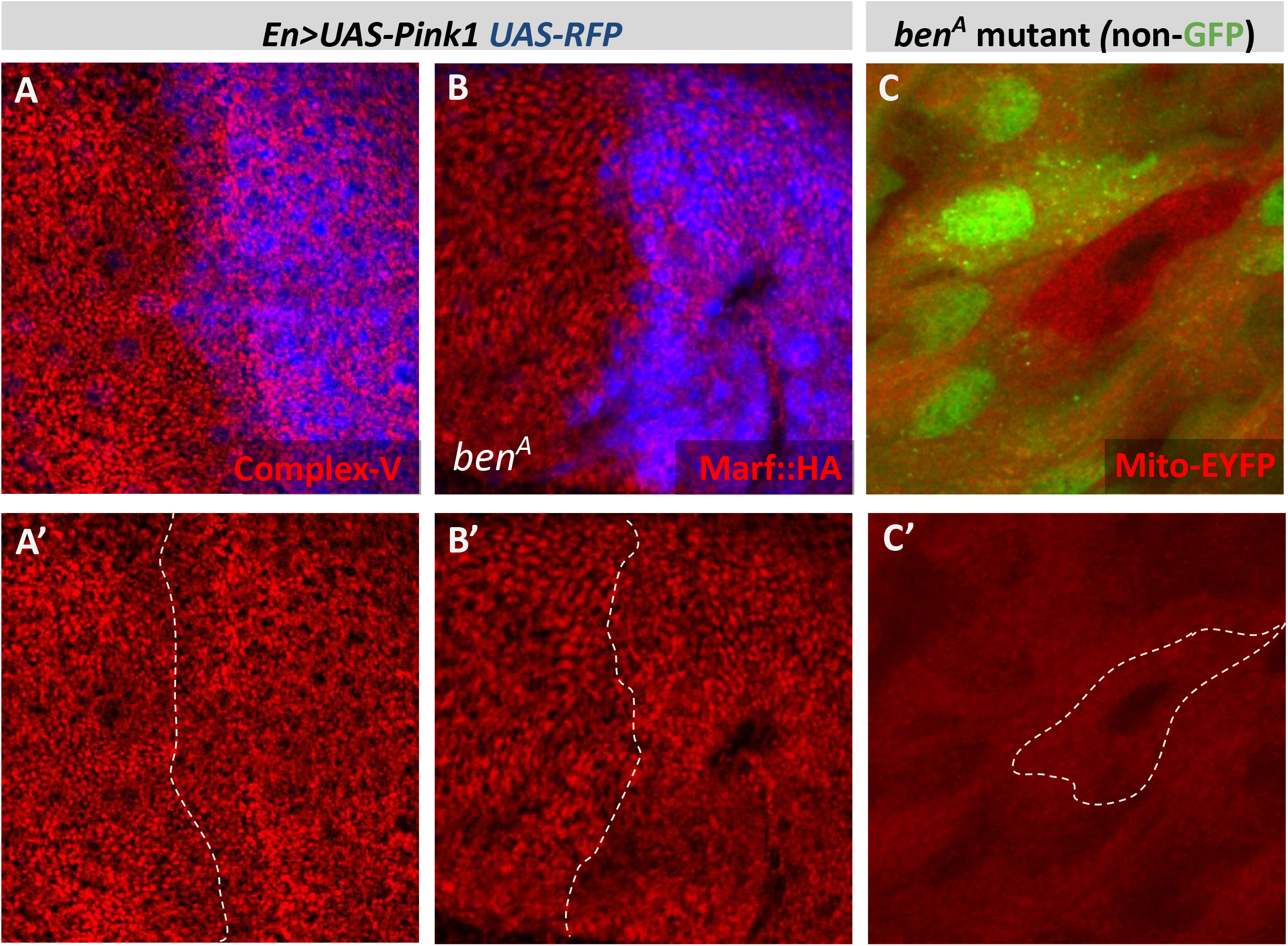
*(A-A*’)Overexpression of *Pink1* using *En*>*Gal4*, wing discs marked with *UAS-RFP* (blue) and immunostained for Complex-V(red). *(B-B*’*)ben*^*A*^ mutant on overexpression of *Pink1* using *En*>*Gal4*, wing discs marked with *UAS-RFP* (blue) and immunostained for Marf::HA (red). *(C-C*’*)ben*^*A*^ mutant clone (non-green cell, C and dashed white line C’), pupal gut 2h APF expressing *Sq>mito-EYFP* (red) shows complete clearance of mitochondria.

**Supplementary Figure 6:**
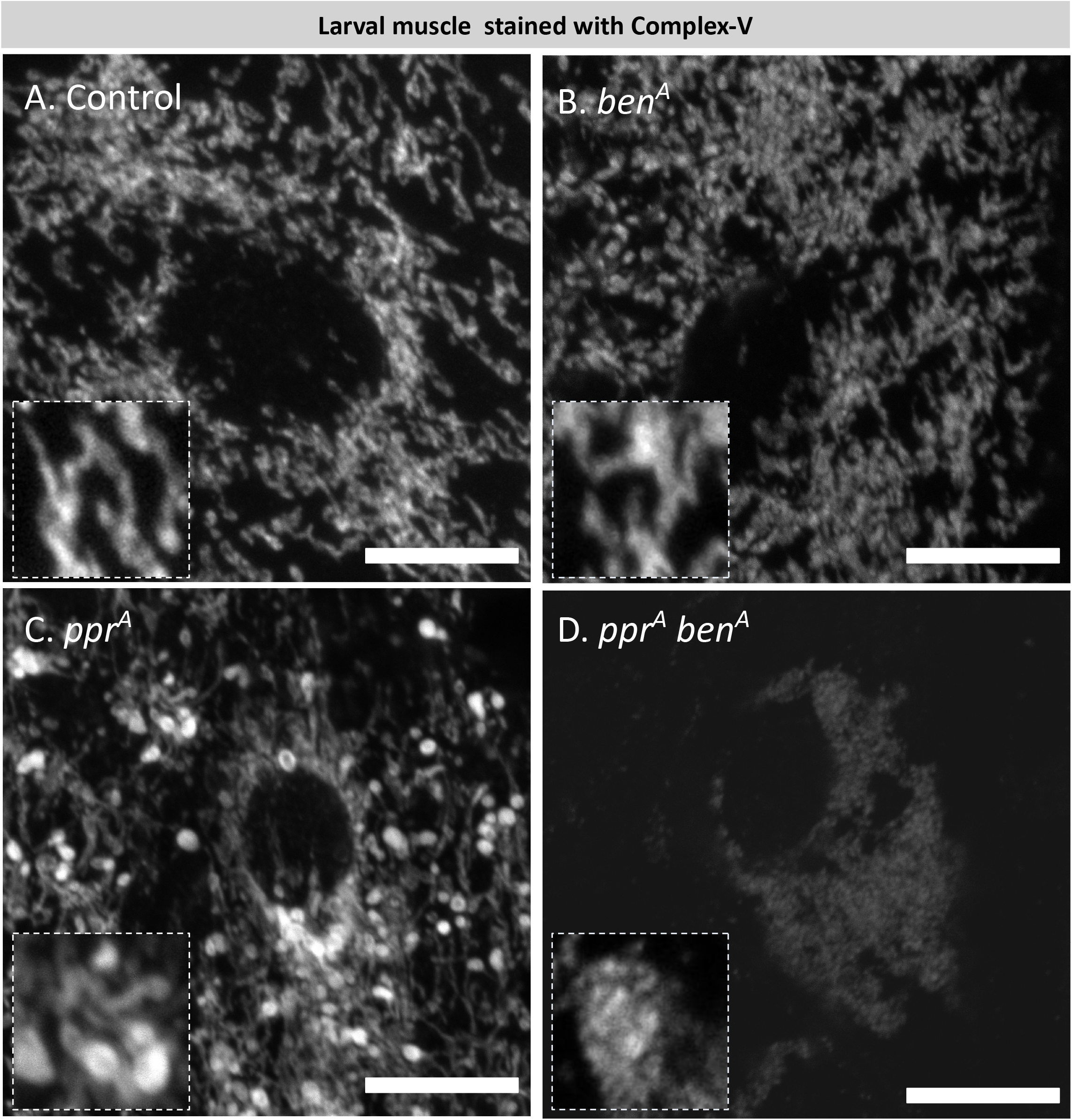
*(A-D)*Confocal sections of third instar larval muscles immunostained for endogenous Complex-V(gray) in control*(A), ben*^*A*^*(B), ppr*^*A*^*(C)* and *ppr*^*A*^ *ben*^*A*^*(D)* larvae.

## Material and Methods

### *Drosophila* culture

Flies were cultured on standard media containing sucrose, malt, yeast and corn flour at room temperature. Crosses were maintained at 25°C. Crosses involving RNAi were maintained at 28°C. *Drosophila* larvae expressing *UAS-Prosβ6*^*1*^ were maintained at 25°C till 3^rd^ instar stage, and were then transferred to 28°C for 24 hours before dissection, to avoid cell death observed on prolonged inhibition of proteasomal activity. To activate the *FLP*-FRT system, heat shock was given during first instar larval stages at 37°C for 1hr. Genotypes used are as listed in Table 1. For drug treatments 3^rd^ instar larvae were transferred to food containing 3mM chloroquine, 100µM MG132, or DMSO (vehicle control) for 24 hours prior to dissection. For western blot and qPCR, 3^rd^ instar larvae were used. We observed that development of *ppr*^*A*^ mutant larvae is substantially delayed. Therefore, we used size matched 3^rd^ instar *ppr*^*A*^ mutant larvae that are obtained after 14-15 days post hatching.

**TABLE 1.**

| **Genotype** | **Source** |
| --- | --- |
| **Figure 1** |  |
| *y<sup>l</sup> ppr<sup>A</sup> w\* P{neoFRT}19A* | FBst0067166<br>(29) |
| *Marf::HA* (Genomic tag on Chromosome III) | (32) |
| *Ubiquitin>GFP hsFLP [122], P{neoFRT}19A* | Hugo Bellen |
| *Ubiquitin>GFP, hsFLP [122], P{neoFRT}19A ;;*<br>*Marf::HA(III)* | This study |
| *Tom20-mCherry* (Genomic construct Chromosome III) | (95) |
| *Ubiquitin>GFP, hsFLP [122], P{neoFRT}19A;;*<br>*Tom20-mCherry* | This study |
| *w;; pacman Opal::HA::Flag VK31 (III) (Opal::HA)* | This study |
| *Ubiquitin>GFP, hsFLP [122], P{neoFRT}19A ;;*<br>*Opal::HA::Flag VK31* | This study |
| *y<sup>l</sup> w\* ; P{w[+mC]=FLAG-FlAsH-HA-Drp1}3, Ki[1]* | FBst0042208<br>(96) |
| *Ubiquitin>GFP, hsFLP [122], P{neoFRT}19A ;;*<br>*P{w[+mC]=FLAG-FlAsH-HA-Drp1}3, Ki[1]* | This study |
| *y<sup>l</sup> w\* P{neoFRT}19A* | (30) |
| **Figure 2** |  |
| *y<sup>l</sup> ppr<sup>A</sup> w\* P{neoFRT}19A* | FBst0067166<br>(29) |
| *w ;; Actin>Gal4* | Hugo Bellen |
| *w ;; Actin>Gal4, Marf::HA (III)* | This study |
| *y<sup>l</sup> ppr<sup>A</sup> w\* P{neoFRT}19A;; Actin>Gal4, Marf::HA (III)* | This study |
| *w\*; P{UAS-Prosbeta6[1].B}2B (II)* | FBst0006786<br>(42) |
| *Ubiquitin>GFP, hsFLP P{neoFRT}19A*<br>*;P{UAS-Prosbeta6[1].B}2B (II)* | This study |
| **Figure 3** |  |
| *y<sup>l</sup> ppr<sup>A</sup> w\* P{neoFRT}19A* | FBst0067166<br>(29) |
| *w\* PinkI<sup>5</sup>* | FBst0051649<br>(48) |
| *y<sup>l</sup> ppr<sup>A</sup> PinkI<sup>5</sup> P{neoFRT}19A* | This study |
| *w\*;; park<sup>A21</sup> (III)* | FBst0051652 |
|  | (97) |
| $y^l ppr^A w^* P\{neoFRT\}19A;; park^{A21} Marf::HA$ | This study |
| *Ubiquitin>GFP, hsFLP [122], P{neoFRT}19A ;; park<sup>A21</sup>* | This study |
| *Ubiquitin>GFP, hsFLP [122], P{neoFRT}19A;*<br>*P{Marf::HA}2* | This study |
| **Figure 4** |  |
| $y^l w^* ben^A P\{neoFRT\}19A$ | FBst0057057<br>(30) |
| *Ubiquitin>GFP, hsFLP [122], P{neoFRT}19A ;;*<br>*Marf::HA (III)* | This study |
| *Ubiquitin>GFP, hsFLP [122], P{neoFRT}19A ;;*<br>*Tom20::mCherry (III)* | This study |
| $w; P\{Marf::HA\}2 P\{w[+mW.hs]=en2.4-GAL4\}e16E,$<br>$P\{w[+mC]=UAS-RFP.W\}2$ | This study |
| $w^*;;P\{UAS.ben::V5.w+mC\}$ | This study |
| $y^l ppr^A w^* P\{neoFRT\}19A$ | FBst0067166<br>(30) |
| $y^l ppr^A w^* ben^A P\{neoFRT\}19A$ | This study |
| **Figure 5** |  |
| $w; P\{Marf::HA\}^2 P\{w[+mW.hs]=en2.4-GAL4\}e16E,$<br>$P\{w[+mC]=UAS-RFP.W\}^2$ | This study |
| $w^*; P\{w[+mC]=UAS-Pink1.C\}A$ | FBst0051648<br>(48) |
| $y^l w^* ben^A P\{neoFRT\}19A ; P\{Marf::HA\}^2$<br>$P\{w[+mW.hs]=en2.4-GAL4\}e16E,$<br>$P\{w[+mC]=UAS-RFP.W\}^2$ | This study |
| $Ubiquitin>GFP, hsFLP$<br>$[122], P\{neoFRT\}19A;; P\{w[+mC]=UAS-Pink1.C\}A$ | This study |
| $P\{Pink1-9Myc\}$ (Pink1::Myc) (II) | FBtp0022940<br>(48) |
| $y^l w^* ben^A P\{neoFRT\}19A; P\{Pink1-9Myc\}$ | This study |
| $y^l w^* P\{neoFRT\}19A; P\{Pink1-9Myc\}$ | This study |
| **Figure 6** |  |
| $y^l w^* P\{neoFRT\}19A$ | (30) |
| $y^l w^* ben^A P\{neoFRT\}19A$ | FBst0057057<br>(30) |
| $y^l ppr^A w^* P\{neoFRT\}19A$ | FBst0067166<br>(29) |
| $y^l ppr^A ben^A P\{neoFRT\}19A$ | This study |
| *cl<sup>l</sup> w\* FRT19A/ Dp(1;Y)y+ v+; ey-FLP* | (30) |
| **Supplementary Figure 1** |  |
| *y<sup>l</sup> ppr<sup>A</sup> w\* P{neoFRT}19A* | FBst0067166<br>(29) |
| *y<sup>l</sup> ppr<sup>E</sup> w\* P{neoFRT}19A* | FBst0067167<br>(29) |
| *P{Ubi-mRFP.nls}, w\* P{hsFLP}, P{neoFRT}19A* | FBst0031418 |
| *w\*;P{w[+mC]=Marf-gHA}2* (Genomic construct Chromosome II) | FBst0067156<br>(32) |
| *P{Ubi-mRFP.nls}, w\* P{hsFLP}, P{neoFRT}19A; P{Marf-gHA}2* | This study |
| *Marf::mCherry* (Genomic rescue construct, Chromosome III) | Hugo Bellen |
| *Ubiquitin>GFP, hsFLP [122], P{neoFRT}19A ;; Marf::mCherry* (III) | This study |
| *y<sup>l</sup> w\* P{neoFRT}19A* | (30) |
| *Ubiquitin>GFP hsFLP [122], P{neoFRT}19A* | Hugo Bellen |
| **Supplementary Figure 2** |  |
| *y<sup>l</sup> w\* HUWE1<sup>B</sup> P{neoFRT}19A* | FBst0052343 |
|  | (30) |
| *ppr<sup>A</sup> HUWE1<sup>B</sup> P{neoFRT}19A* | This study |
| *Ubiquitin>GFP, hsFLP [122], P{neoFRT}19A ;;*<br>*Marf::HA (III)* | This study |
| *MUL1<sup>A6</sup> (III)* | FBal0301081<br>(47) |
| *y<sup>l</sup> ppr<sup>A</sup> w\* P{neoFRT}19A ;; MUL1<sup>A6</sup>* | This study |
| *Ubiquitin >GFP, hsFLP [122] ,P{neoFRT}19A ;;*<br>*MUL1<sup>A6</sup>* | This study |
| *y<sup>l</sup> w\* P{neoFRT}19A* | (30) |
| *w<sup>1118</sup>;P{w[+mW.hs]=en2.4-GAL4}e16E,*<br>*P{w[+mC]=UAS-RFP.W}2* | FBst0030577 |
| *w; P{Marf::HA}2 P{w[+mW.hs]=en2.4-GAL4}e16E,*<br>*P{w[+mC]=UAS-RFP.W}2* | This study |
| *y<sup>l</sup> ppr<sup>A</sup> w\* P{neoFRT}19A;P{Marf::HA}2*<br>*P{w[+mW.hs]=en2.4-GAL4}e16E,*<br>*P{w[+mC]=UAS-RFP.W}2* | This study |
| *y<sup>l</sup> v<sup>l</sup>; P{y[+t7.7] v[+t1.8]=TRiP.JF02007}attP2 (RNAi*<br>*against crc)* | FBst0025985 |
| *Ubiquitin >GFP, hsFLP [122], P{neoFRT}19A;;*<br>*P{y[+t7.7] v[+t1.8]=TRiP.JF02007}attP2* (RNAi against *crc*) | This study |
| *w<sup>1118</sup>; P{GD1425}v3781* (RNAi against *dve*) | FBti0084290 |
| *Ubiquitin >GFP, hsFLP [122], P{neoFRT}19A;;*<br>*P{GD1425}v3781* (RNAi against *dve*) | This study |
| *y<sup>l</sup> v<sup>l</sup>; P{y[+t7.7] v[+t1.8]=TRiP.JF02734}attP2* (RNAi against *foxo*) | FBst0027656 |
| *Ubiquitin >GFP, hsFLP [122], P{neoFRT}19A;;*<br>*P{y[+t7.7] v[+t1.8]=TRiP.JF02734}attP2* (RNAi against *foxo*) | This study |
| *P{UAS-rOTC.P} (UAS-OTC)* | FBal0291051<br>(56) |
| *P{UAS-rOTC.d} (UAS-ΔOTC)* | FBal0291052<br>(56) |
| **Supplementary Figure 3** |  |
| *P{FRT(w<sup>hs</sup>)}2A* | FBst0001997 |
| *park<sup>Δ21</sup> P{FRT(w<sup>hs</sup>)}2A* | This study |
| *P{hsFLP}1, y1 w\* P{UAS-mCD8::GFP.L}Ptp4ELL4;*<br>*P{tubP-GAL80}LL9 P{FRT(whs)}2A* | FBst0044404 |
| *Pink1<sup>5</sup> P{neoFRT}19A* | This study |
| *Ubiquitin>GFP, hsFLP [122], P{neoFRT}19A ; ;*<br>*Marf::HA (III)* | This study |
| *y<sup>l</sup> w<sup>\*</sup> ben<sup>B</sup> P{neoFRT}19A* | FBst0057058<br>(30) |
| *P{Ubi-mRFP.nls}, w<sup>\*</sup> P{hsFLP}, P{neoFRT}19A ;*<br>*P{Marf::HA}2* | This study |
| *y<sup>l</sup> ppr<sup>A</sup> w<sup>\*</sup> ben<sup>B</sup> P{neoFRT}19A* | This study |
| *w<sup>\*</sup> ; P{UAS-HA::ben.w+mC}* | This study |
| *y<sup>l</sup> w<sup>\*</sup> ; ; P{w[+mC]=tubP-GAL4}LL7 (III)* | FBti0012687 |
| *w<sup>\*</sup> ; ; P{UAS.ben::V5.w+mC}* | This study |
| **Supplementary Fig 4** |  |
| *w<sup>\*</sup> ; P{w[+mC]=UAS-Pink1.C}A* | FBst0051648<br>(48) |
| *w ; P{Marf::HA}2 P{w[+mW.hs]=en2.4-GAL4}e16E,*<br>*P{w[+mC]=UAS-RFP.W}2* | This study |
| *y<sup>l</sup> w<sup>\*</sup> ben<sup>A</sup> P{neoFRT}19A ; P{Marf::HA}2*<br>*P{w[+mW.hs]=en2.4-GAL4}e16E,*<br>*P{w[+mC]=UAS-RFP.W}2* | This study |
| *P{Pink1-9Myc} (Pink1::Myc)* | FBtp0022940<br>(48) |
| *y<sup>l</sup> w<sup>\*</sup> ben<sup>A</sup> P{neoFRT}19A; P{Pink1-9Myc}* | This study |
| *y<sup>l</sup> w<sup>\*</sup> P{neoFRT}19A; P{Pink1-9Myc}* | This study |
| *w<sup>\*</sup>; P{sqh-EYFP-Mito}(III)* | FBst0007194 |
| *P{Ubi-mRFP.nls}, w<sup>\*</sup> P{hsFLP}, P{neoFRT}19A;;*<br>*P{sqh-EYFP-Mito}(III)* | This study |
| **Supplementary Figure 5** |  |
| *y<sup>l</sup> w<sup>\*</sup> P{neoFRT}19A* | (30) |
| *y<sup>l</sup> w<sup>\*</sup> ben<sup>A</sup> P{neoFRT}19A* | FBst0057057 |
| *y<sup>l</sup> ppr<sup>A</sup> w<sup>\*</sup> P{neoFRT}19A* | FBst0067166<br>(29) |
| *y<sup>l</sup> ppr<sup>A</sup> w<sup>\*</sup> ben<sup>A</sup> P{neoFRT}19A* | This study |

### Generation of transgenic flies

*ben* sequence was amplified from genomic DNA. These PCR amplified *ben* ORF sequences were then inserted into a pUAST vector containing attB sites, flanking the insert using EcoRI-XhoI. pUAST vectors containing *UAS-ben::V5/UAS-HA::ben* were injected into embryos containing attP2 landing site and integrase. Transgenic flies were selected based on presence of w^+mC^. Primers used: P{UAS.*ben::V5*.w^+mC^}: Fwd-5’-GGAATTCGCCACCATGTCCAGCCTGCCACGTC-3’ and Rev-5’-CCGCTCGAGTTACGTAGAATCGAGACCGAGGAGAGGGTTAGGGATAGGCTTACCGTCTTCGACGGCATAT-3’. P{UAS-*HA::ben*.w^+mC^}: Fwd-5’-GGAATTCGCCACCATGTACCCATACGACGTCCCAGACTACGCTATGTCCAGCCTGCCACGTC-3’ and Rev-5’-CCGCTCGAGTCAGTCTTCGACGGCATAT-3’.

Opa1::3FLAG-2HA genomic construct was generated using the P(acman) system (93). Briefly, the 3FLAG-2HA tag was amplified from C-terminal tag fusion vector pL452-C-3FLAG-2HA and inserted at the C terminal of Opa1 through recombineering in the P(acman) clone CH322-27B08, which was subsequently injected into *y*^*1*^ *w*^*1118*^ ; PBac{y+-attP-3B}VK00033 flies.

### Immunofluorescence and imaging

Larvae were dissected in 1X PBS, followed by fixing in 4% paraformaldehyde (Himedia - TCL-119 - 100ml) for 30 minutes at room temperature and three washes in 1X PBS with 0.2% TritonX-100 (Himedia - MB031, 1X PBST). Primary antibodies were incubated overnight at 4°C. Followed by blocking in 5% normal goat serum (Himedia - RM10701) for 1h at room temperature and then secondary antibody incubation followed by washing and dissection. Samples were mounted in Vectashield (VectorLabs - H100) and imaged under 40X or 63X oil immersion Leica Stellaris 5 or Olympus FV3000 confocal microscopes. Images were processed using Fiji. All antibody dilutions and the blocking solution were made in 1X PBST; details of antibodies and their dilutions used are listed in Table 2.

**TABLE 2.**

| **Antibody** | **Catalog** | **Dilution** |
| --- | --- | --- |
| Mouse HA | Cell Signaling Technology- 2367S | 1:500 IF |
| Rabbit HA | Cell Signaling Technology- 3724S | 1:500 IF |
| Mouse Complex-V | Abcam- 176569 | 1:500 IF<br>1:2500 WB |
| Rabbit V5 | Cell Signaling Technology- 13202S | 1:500 IF |
|  |  | 1: 5000 WB |
| Rabbit mCherry | Cell Signaling Technology- 43590S | 1:500 IF |
| Rabbit Actin | Cell Signaling Technology- 4967S | 1:5000 WB |
| Mouse Actin | Invitrogen- MA5- 15739 | 1:5000 WB |
| Rabbit Tubulin | Novus Biologicals- NB100-56459 | 1:1000 WB |
| Rabbit Hsp60A | (98) | 1:200 IF |
| Mouse Myc Tag | Cell Signaling Technology-2276S | 1:1000 WB |
| Rabbit p62 | Abcam- ab178440 | 1:500 IF |
| Anti-Mouse 488 | Invitrogen- A11029 | 1:500 IF |
| Anti Mouse 555 | Invitrogen- A21424 | 1:500 IF |
| Anti-Mouse 633 | Invitrogen- A21052 | 1:500 IF |
| Anti-Rabbit 488 | Invitrogen- A32731 | 1:500 IF |
| Anti-Rabbit 555 | Invitrogen- A32732 | 1:500 IF |
| Anti-Rabbit 647 | Invitrogen- A32733 | 1:500 IF |
| Anti-Rabbit HRP | Novus Biologicals- NB7160 | 1:5000 WB |
| Anti-Mouse HRP | Novus Biologicals- NB7539 | 1:5000 WB |

IF- Immunofluorescence WB- Western Blot

### Eye phenotype imaging

Mutant eyes were created by crossing heterozygous mutant flies with *w cl(1)* FRT19A /Dp(1;Y); *ey-FLP* flies. The eye images were then acquired on a Leica M205FA Stereo Zoom microscope.

### Mitochondrial fractionation

Thirty 3^rd^ instar larvae were collected and washed with chilled mitochondrial isolation buffer (210mM mannitol, 70mM sucrose, 1mM EGTA, 5mM HEPES, and 0.5% BSA). The larvae were then transferred into a 1.5ml centrifuge tube containing 250µl of chilled mitochondrial isolation buffer. Micro-pestles were used to homogenize the samples, keeping the sample on ice. 300 µl of mitochondrial isolation buffer was added to the lysate. Lysate was centrifuged at 200G for 5 mins at 4°C to remove large debris. The supernatant was then centrifuged at 1500G for 5mins at 4°C and the pellet was discarded. The supernatant was centrifuged at 8000G for 15mins at 4°C.

The supernatant containing the cytoplasmic fraction was processed for western blot by adding Laemmli buffer (0.004% bromophenol blue, 20% glycerol, 4% SDS and 0.125M Tris-HCl pH 6.8) having 5% beta-mercaptoethanol and heated at 98°C for 5mins. The pellet was resuspended in 550µl fresh mitochondrial isolation buffer and again centrifuged at 8000G for 15 mins at 4°C. The pellet containing the mitochondrial fraction was resuspended in 100µl of Laemmli buffer having 5% beta-mercaptoethanol and heated at 98°C for 5mins.

### Western blot

3^rd^ instar larvae were crushed in RIPA lysis buffer [50mM Tris,150mM NaCl, 0.2% Triton X 100 and 1X protease and phosphatase inhibitor cocktail (Thermo Fisher - A32965,A32957 respectively)], followed by centrifugation at 16,000g for 10 mins at 4°C. Clear fat free supernatant was used for total protein estimation. Lysate was mixed with equal volume of 1X Laemmli buffer (0.004% bromophenol blue, 20% glycerol, 4% SDS and 0.125M Tris-HCl pH 6.8) having 5% beta-mercaptoethanol and heated at 98°C for fine minutes, centrifuged, and 25µg of protein was loaded in each well and resolved on 4-15% gradient Tris-Glycine gel (Bio-Rad - 4561086). Semi-dry transfer was done onto 0.2µm Nitrocellulose membrane as per Trans-BlotTurbo Kit (Bio-Rad - 1704270) for seven minutes. Blocking in either 5% Blotto (Santa Cruz sc - 2325) or 5% BSA made in 1X TBS with 0.1% Tween-20 (1X TBSTw20) for 1 hour at room temperature followed by primary antibody incubation overnight at 4°C. After washing thrice in 1X TBSTw20, membranes were incubated in HRP conjugated secondary antibodies (Table2) for 2 hours at room temperature. After washing, they were developed using Clarity Western ECL Substrate (Bio-Rad - 1705061) and visualized using Vilber-Lourmat chemidoc. Band intensities were quantified using Fiji and normalized with βActin.

### Real-time PCR

3^rd^ instar larvae were used for RNA isolation using TRIzol (Ambion life tech - 15596018) method. cDNA conversion for 1µg of RNA was carried out using a cDNA conversion kit (Thermo Fisher - 4368814). qPCR was carried out in 96 well plates in three technical replicates for each of the three biological replicates. *Marf* qPCR was done using the iTaq SYBR Green supermix (Bio-Rad -1725121) using LightCycler 96 (Fig. 5F).

Following primers were used :

Marf-Fwd-5’-CGAGTGCCAGGAATCGGTTA-3’, Marf-Rev5’-ATCTGAAAGCCCTCGGCAAT-3’, RP49-Fwd-5’-TCCTACCAGCTTCAAGATGAC-3’, RP49-Rev-5’-CACGTTGTGCACCAGGAACT-3’.

### TMRE Staining

3^rd^ instar larvae were dissected in Schneider’s Insect media (Himedia - IML003-500ml). The larvae were transferred to media containing 100nM TMRE (Thermo Fisher - T669) in Schneider’s Insect media and incubated for 20 mins. The wing discs were dissected and mounted in Schneider’s media using a coverslip. The tissues were live imaged using Leica Stellaris 5 confocal microscope at 63X oil objective.

### Mitochondrial morphology analysis

Wing discs immunostained for Complex-V were imaged using Leica Stellaris 5 confocal microscope at 63X oil objective. The mitochondria were segmented on Fiji using the Trainable Weka segmentation plugin (94). The segmented images were then used to find out mitochondrial area using Particle Analyze Tool on Fiji.

Blind test: For qualitative assessment of mitochondrial morphology in larval muscle, we renamed a set of images containing mitochondria from larval muscles with random numbers. The images from different genotypes (*control, ben*^*A*^, *ppr*^*A*^, and *ppr*^*A*^ *ben*^*A*^) were pooled and were assessed for the presence of different mitochondrial morphologies, including presence or absence of mitochondria network, large globular mitochondria, ring shaped mitochondria and mitochondrial aggregates. Multiple images were used for the assessment, 40 images from 11 larvae for control, 24 images from 7 larvae for *ben*^*A*^, 27 images from 7 larvae for *ppr*^*A*^, and 32 images from 9 larvae for *ppr*^*A*^*ben*^*A*^.

### Statistics analysis

At least three independent experiments were used for all quantifications, the n values for each experiment is indicated in their respective figure legends. Two-tailed paired t-test was used to analyze data obtained from clonal analysis, One sample t-test was used to analyze the data in Supplementary Fig. 5. Two-tailed unpaired t-test was used to analyze all other data sets. Significance of the data was represented as * for p<0.05, ** for p<0.01, and *** for p<0.0001. Details of the test used and the significance is mentioned in respective figure legends.

## Acknowledgements

We thank Hong Xu, Ming Guo, Luis Martins, Samantha Loh and Hugo J Bellen for the various fly lines used in this work. Stocks obtained from the Vienna Drosophila Resource Center and Bloomington Drosophila Stock Center (NIH P40OD018537) were used in this study. We thank NCBS-TIFR, Banglore fly facility for embryo injections. MJ is supported by the Department of Atomic Energy (Project Identification No. RTI 4007), Department of Science and Technology, SERB (CRG/2020/003275), Department of Biotechnology (BT/PR32873/BRB/10/1850/2020), Government of India. MJ is a Ramalingaswami fellow, Department of Biotechnology, Government of India, under project number BT/RLF/Re-entry/06/2016. RNC and TA are supported by intramural funding of TIFR. SNJ is supported by DBT/Wellcome trust India Alliance (grant no IA/I/18/1/503629) and intramural funding of CSIR-Centre for Cellular and Molecular Biology, Hyderabad, India. We thank Titus P Ponrathnam for cloning of Ben::V5 construct preparation. We acknowledge Aravind H and Sayantan Datta for critical reading and editing of the manuscript and the MJ lab members for fruitful discussion.

